# Identification and Evaluation of Benzimidazole-Agonists of Innate Immune Receptor NOD2

**DOI:** 10.1101/2025.07.03.661155

**Authors:** Liora Wittle, Karl Ocius, Mahendra Chordia, Carly van Wagoner, Timothy Bullock, Marcos M. Pires

**Affiliations:** Department of Chemistry University of Virginia, Charlottesville, VA, United States 22904; Department of Pathology University of Virginia Charlottesville, VA, United States 22908

## Abstract

Emerging evidence has demonstrated the importance of pattern recognition receptors (PRRs), including the nucleotide-binding and oligomerization domain receptor 2 (NOD2), in human health and disease states. NOD2 activation has shown promise with aiding malnutrition recovery, lessening irritable bowel disease (IBD) symptoms, and increasing the efficacy of cancer immunotherapy. Currently, most NOD2 agonists are derivatives or analogs of the endogenous agonist derived from bacterial peptidoglycan, muramyl dipeptide (MDP). These MDP-based agonists can suffer from low oral bioavailability and cause significant adverse side effects. With the goal of broadly improving NOD2 therapeutic interventions, we sought to discover a novel small molecule capable of activating NOD2 by screening a library of total 1917 FDA approved drugs in a phenotypic assay. We identified a class of compounds, benzimidazoles, that act as NOD2 agonists, with the most potent member of this class being nocodazole. Nocodazole activates NOD2 with nanomolar potency and causes the release of cytokines canonically associated with MDP-induced NOD2 activation, suggesting its potential to elicit similar therapeutic immune effects as MDP and potentially offer improved pharmacological properties.

## INTRODUCTION

Nucleotide binding and oligomerization domain receptor 2 (NOD2) is a pattern recognition receptor (PRR) expressed mainly on immune and gut epithelial cells.^1–5^ PRRs detect pathogen associated molecular patterns (PAMPs) and activate the immune system against potential infections.^6^ PRRs are increasingly targeted for treating or preventing infections and inflammatory diseases.^5,7,8^ NOD2 – a cytosolic, membrane associated receptor – detects fragments of peptidoglycan (PG), a primary component of bacterial cell membranes expressed by all known species of bacteria, including those associated with the gut microbiome.^4,5^ The structure of PG consists of crosslinked units of *N*-acetyl glucosamine (GlcNAc) and *N*-acetyl muramic acid (MurNAc), with a short peptide chain conjugated to MurNAc.^9^ Muramyl dipeptide (MDP), the minimal peptidoglycan fragment capable of activating NOD2, does so at nanomolar concentrations.^10^ Upon activation, NOD2 initiates signaling through receptor interacting serine/threonine kinase 2 (RIPK2), leading to the activation of nuclear factor kappa beta (NF-kB) (**Fig. 1a**). NF-kB activation, in turn, causes the transcription of several cytokines and other immune signaling molecules, such as interleukin (IL)-6, IL-8, IL-10, and tumor necrosis factor alpha (TNF-α) that can trigger and modulate inflammatory responses.^5,11,12^ Despite the promising features of MDP, such as its potency in activating NOD2, early clinical evaluations revealed it to be pyrogenic and to have poor oral bioavailablity.^13^

**Figure 1.**
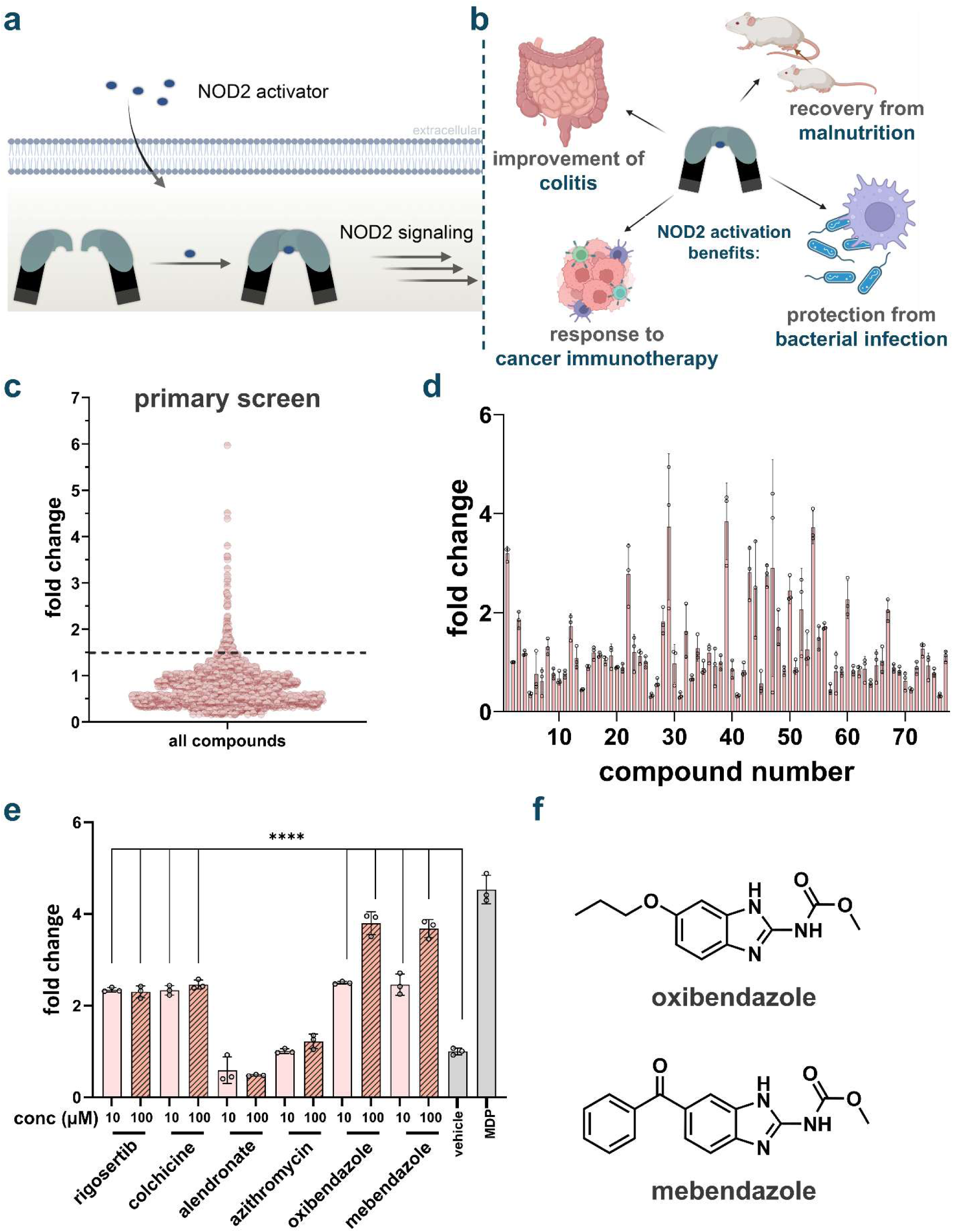
**(a)** NOD2 is an intracellular pattern recognition receptor that becomes activated upon sensing muramyl dipeptide (MDP), a conserved motif from bacterial peptidoglycan. Upon activation, NOD2 oligomerizes and recruits downstream signaling molecules, such as RIPK2, leading to activation of NF-κB and MAPK pathways and subsequent inflammatory cytokine production. **(b)** Physiological functions that have been associated with NOD2 activation. **(c)** Screening of FDA approved drug library for NOD2 activation. Initial screening of compounds was performed in HEK-Blue cells, which were treated with each compound at 100 µM in DMSO for 16 h (n=1). The analysis was performed by measuring the absorbance of each well at 655 nm. Fold change calculated by compound absorbance/average background. **(d)** Secondary screening of potential hits from the primary screen of FDA-approved compounds for NOD2 activation. HEK-Blue cells were treated with each compound at 100 µM in DMSO for 16 h using a colorimetric assay (n = 3. **(e)** HEK-Blue cells were treated with each compound at 10 µM and 100 µM for 16 h using a colorimetric assay (n=3). P values determined by one-way ANOVA (**** p< 0.0001). Unlabeled bars have no significance compared to background. **(f)** Chemical structures of oxibendazole and mebendazole.

NOD2 activation has been shown to improve recovery from malnutrition,^14^ regulate appetite,^15^ protect against bacterial infections,^16,17^ reduce inflammatory bowel disease (IBD) symptoms,^4,18^ and enhance efficacy of PD-1/PD-L1 cancer immunotherapy **(Fig 1b)**.^19–21^ Many of the beneficial effects from NOD2 activation have been demonstrated using probiotics or bacterial-derived molecules (e.g., isolated complete PG). For example, strains that release PG-digesting hydrolases in the gut have been shown to improve NOD2 activation by promoting the depolymerization of PG into NOD2-activating fragments.^16,22^ Treatment with bacteria that secrete PG hydrolases, or with isolated hydrolases themselves, has been shown to alleviate IBD symptoms and protect against bacterial infection.^16,22,23^ Another promising therapeutic strategy involves the use of NOD2 agonists as adjuvants in cancer checkpoint therapy. Programmed cell death protein 1 (PD-1) is a receptor, widely expressed on immune cells, that recognizes programmed cell death ligand 1 (PD-L1) and subsequently suppresses immune activity.^24^ Many tumors upregulate PD-L1 to evade immune surveillance by preventing immune cells from recognizing and eliminating cancer cells.^24^ To date, clinical responses to checkpoint therapy have been highly variable across cancer patients.^25^ Notably, stronger responses to PD-1/PD-L1 immunotherapy have been associated with higher levels of NOD2 activation.^19–21,25,26^ In mice, treatment with a probiotic bacterium engineered to express a PG hydrolase enhanced the efficacy of PD-L1 immunotherapy, and this effect was shown to be dependent on NOD2 activation.^19^

While the use of probiotic bacteria to modulate the gut microbiome is promising, it presents several challenges. Probiotics can be difficult to permanently introduce into the host, and if harbored within the gut, their growth rates and release of peptidoglycan (PG) fragments can be challenging to control. Sustaining their residence in the gut is also problematic during dysbiosis, and interactions with the existing microbial community can further complicate outcomes.^27^ In immune compromised patients, probiotic bacteria may also pose additional challenges, as weaker immune systems struggle to control bacterial growth.^28^ To address the limitations of probiotics, alternative strategies have focused primarily on MDP analogs. These efforts include modifying the sugar moiety of MDP or altering the peptide side chains.^29–31^ Compounds such as *N*-arylpyrazole and desmuramylpeptide agonists have demonstrated promising NOD2 activation profiles.^29–31^ However, they still remain structurally similar to MDP and have not been evaluated for their stability, oral bioavailability, or their safety in humans. Accordingly, there remains a critical need for structurally diverse NOD2 agonists that enable controlled NOD2 activation across a broad spectrum of clinical contexts.

We aimed to identify NOD2 agonists from a diverse library of FDA-approved small molecule drugs. These libraries represent an attractive entry point for drug discovery, as the compounds have already undergone extensive safety and toxicity assessments, thereby substantially lowering the time and cost required for clinical translation.^32^ Additionally, they contain structurally diverse scaffolds that have demonstrated favorable pharmacokinetic properties, including oral bioavailability and metabolic stability – characteristics that are often lacking in natural product-derived agonists like MDP. To identify novel NOD2 agonists, we screened an FDA-approved drug library using NOD2-HEK-Blue NF-κB reporter cell lines. Among the full set of screened molecules, two structurally related hit compounds, each containing a benzimidazole moiety, were identified. Subsequent expansion of this chemical scaffold revealed multiple benzimidazoles capable of activating NOD2. Notably, one benzimidazole derivative not included in the original screen, nocodazole, exhibited nanomolar potency and elicited immune responses comparable to those induced by MDP.

## RESULTS AND DISCUSSION

### Identification of Benzimidazole NOD2 Agonists

First, we screened 1971 FDA approved drugs using the L1021 DrugDiscovery^TM^ library (Apexbio Technology). The assay was initiated by incubating human NOD2-HEK-Blue NF-κB reporter cells (HEK-Blue NOD2 cells) with each compound at a concentration of 100 µM. This specific reporter cell line is engineered to have human NOD2 receptors and to express a secreted alkaline phosphatase (SEAP) downstream of NF-κB upon NOD2 activation.^5^ The presence of SEAP in the cell culture medium dephosphorylates a small molecule dye, resulting in a color change that can be detected using UV-vis spectrophotometry (**Fig. S1**). The gastrointestinal (GI) tract is a key site of NOD2 expression and activation, and also one where orally administered compounds can reach millimolar concentrations prior to systemic absorption and dilution.^33,34^ Accordingly, the primary screen was performed at elevated concentrations (100 µM) to mirror the high local compound levels anticipated in the GI tract following oral administration.^34–37^ Each plate included internal controls: MDP as a positive control and DMSO as a negative (vehicle) control. Fold-change values were calculated relative to these internal references.

The primary screen identified 90 candidate compounds that induced at least a 1.5-fold increase in activation relative to background (**Fig. 1c**). This threshold was chosen to ensure that only compounds with activation levels clearly exceeding background were retained. To narrow down our potential hits, we analyzed the usage and administration methods of each candidate compound, removing those less suitable for the goal of NOD2 activation. We excluded compounds that are not typically administered orally, as they would not encounter NOD2 on the gut epithelium. Additionally, some of the compounds that displayed more than 1.5-fold activation had FDA approval as drugs of last resort with high toxicity profiles, therefore, they were eliminated from further evaluation. The remaining 76 compounds were re-screened in triplicate, and the most potent inducers of NOD2 were selected for further analysis. (**Fig. 1d**). Six of the compounds with the highest levels of NOD2 activation were identified and tested at a lower concentration: rigosertib, colchicine, alendronate sodium, azithromycin, mebendazole, and oxibendazole (**Fig. 1e, Fig S2**). The last round of testing demonstrated that alendronate sodium and azithromycin performed significantly worse than the other final hits. Based on our results and existing knowledge about these compounds, we excluded two from further evaluation: rigosertib, due to a lack of dose-dependent activity (**Fig. S3**), and colchicine, due to its high potential for toxicity.^38^ Instead, we selected mebendazole and oxibendazole for additional downstream evaluation (**Fig. 1f**). These compounds displayed high levels of NOD2 activation in the reporter cell line and showed a strong dose dependence. Of note, both mebendazole and oxibendazole are anti-helminth drugs and possess a relatively high safety profile (standard dosage in humans being 200 mg/day as a tablet through gut), allowing them to be administered in larger doses.^39,40^

### Expansion of Benzimidazole Class for NOD2 Activation

The two compounds with the highest potency and dose dependance, oxibendazole and mebendazole, share remarkable structural similarity. As such, we suspected that other compounds within the benzimidazole class might also be NOD2 agonists. A member of the benzimidazole family, GSK717, was previously developed by GSK as a NOD2 inhibitor.^41,42^ GSK717 has high selectivity towards NOD2 and is structurally dissimilar from oxibendazole or mebendazole outside of the core benzimidazole moeity.^41,42^ It was described to act as a competitive inhibitor, suggesting an ability to interact with the binding site of NOD2.^42^ This interaction with the binding site supports the concept that benzimidazole containing molecules could potentially also act as agonists.

To further explore the benzimidazole pharmacophore, we tested twelve additional commercially available benzimidazole derivatives for their ability to activate NOD2 (**Fig. 2a**). Of the compounds tested, approximately half exhibited NOD2-activating activity, with some inducing stronger activation than the original hits identified in the primary screen (**Fig. 2b**). Many of these benzimidazoles are also anti-helminth drugs, such as albendazole and fenbendazole. In the body, albendazole and fenbendazole are metabolized into their sulfoxide and sulfone forms (ricobendazole/albendazole sulfone and oxfendazole/fenbendazole sulfone respectively) which exert the anthelmintic effects.^43–45^ Interestingly, none of these metabolites display any NOD2 activity. Except for benomyl, all NOD2-activating benzimidazoles were active at both 1 µM and 10 µM, showing at least a two-fold increase over background at the lower concentration. These results suggest that benzimidazoles may represent a new and general class of NOD2-activating compounds. Subsequent testing revealed that two of the fourteen compounds, parbendazole and nocodazole, retained their NOD2 activation at sub-micromolar concentrations (**Fig. 2c**). Further evaluation analysis showed that these two compounds activate NOD2 with EC_50_ values of 116.2 nM and 315.5 nM for parbendazole and nocodazole, respectively; notably, these new compounds both demonstrated a substantial improvement over one of the top hits from the primary screen (**Fig. 2d**). Although MDP induces higher NOD2 activity than both parbendazole and nocodazole, orally administered drugs become concentrated in the gut, occasionally reaching the millimolar range, and NOD2 receptors are present on the intestinal epithelium, allowing for a greater range of clinically relevant concentration for NOD2 agonists.^33–36^ The range in their biological activity as NOD2 activators indicates a specific pharmacophore that is responsible for the activation of NOD2 and suggests that these lead agents have potential for further development as NOD2-driven immunomodulatory agents.

**Figure 2.**
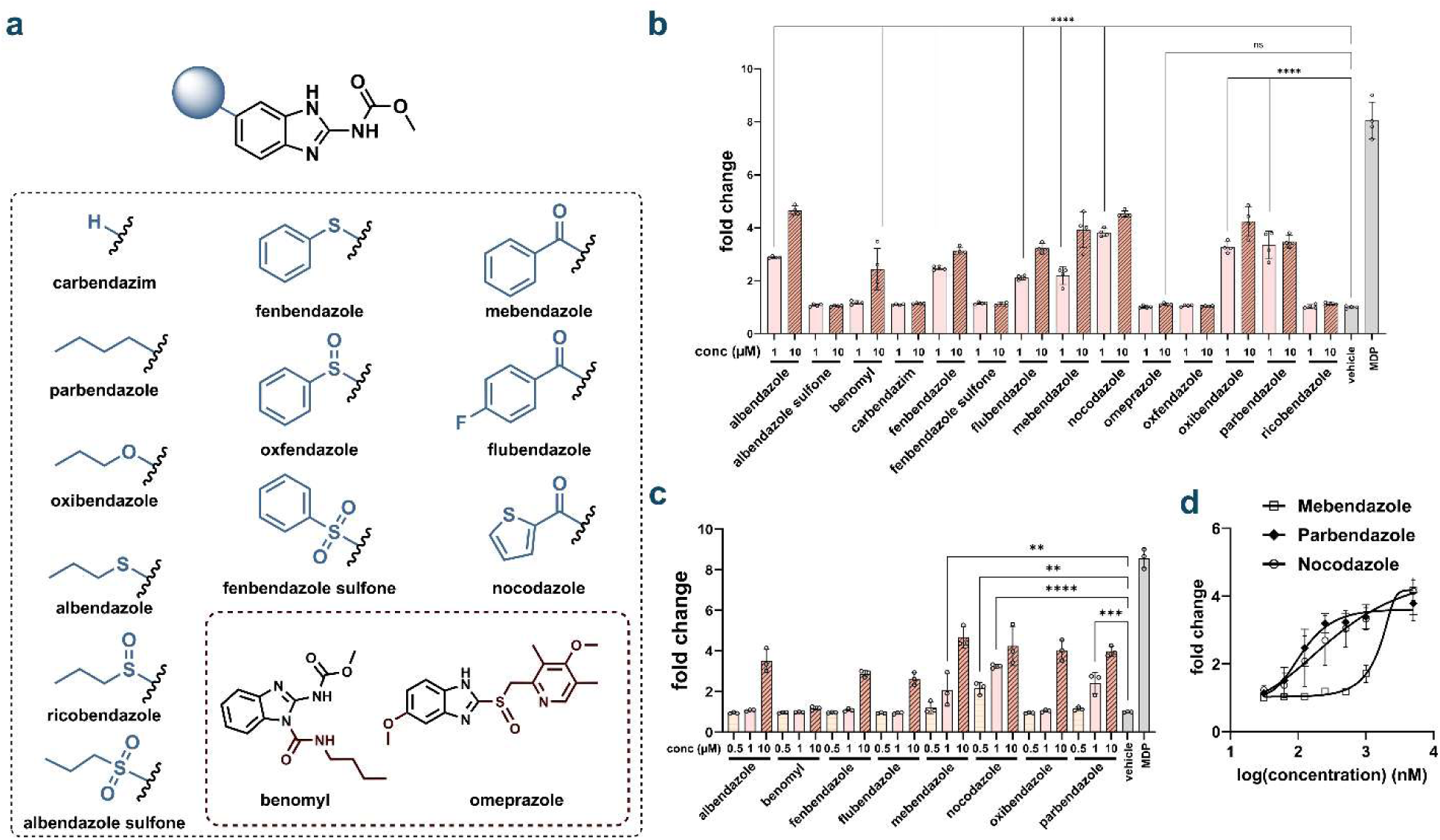
**(a)** Chemical structures of a family of benzimidazole-based compounds that were selected for potential NOD2 activation based on the results from the primary screen. **b)** NOD2 activation of benzimidazole compounds. HEK-Blue cells were treated with each compound at indicated concentrations for 16 h using a colorimetric assay (n=3). The analysis was performed by measuring the absorbance of each well at 655 nm. Fold change calculated by compound absorbance/average background. p values calculated by one-way ANOVA (ns=not significant, **** p<0.0001). Bars mark the lowest concentration with statistically significant activation over background. **(c)** Dose dependence of NOD2 agonist benzimidazole compounds. HEK-Blue cells were treated with each compound at indicated concentrations for 16 h using a colorimetric assay (n=3). **(d)** EC_50_ curves of mebendazole, parbendazole, and nocodazole. HEK-Blue cells were treated with each compound at indicated concentrations for 16 h using a colorimetric assay (n=3). The analysis was performed by measuring the absorbance of each well at 655 nm. Fold change calculated by compound absorbance/average background.

### Confirmation of NOD2-specific Activation by Benzimidazoles

To evaluate the specificity of benzimidazole-mediated NOD2 activation, we assessed cross-reactivity with NOD1 using HEK-Blue NOD1 reporter cell lines. NOD1 is another innate immune receptor that detects PG fragments^2^ with its cognate ligand being γ-D-glutamyl-*meso*-diaminopimelic acid (iE-DAP).^46^ Both NOD1 and NOD2 exhibit remarkable structural homology, sharing caspase-activating and recruitment domains (CARD), nucleotide-binding oligomerization domains (NOD), and leucine-rich repeat regions (LRR) (**Fig. 3a**).^47,48^ This high degree of structural similarity makes NOD1 particularly susceptible to off-target effects from NOD2 agonists, rendering it an appropriate counter-screen for assessing compound selectivity. When evaluated in the NOD1 reporter system, benzimidazoles exhibited only background activation at both 10µM and 1 µM (**Fig. 3b**), demonstrating minimal cross-reactivity. The inability of benzimidazoles to activate NOD1, despite its structural similarity to NOD2, provides strong evidence for the specificity of these compounds as selective NOD2 agonists.

**Figure 3.**
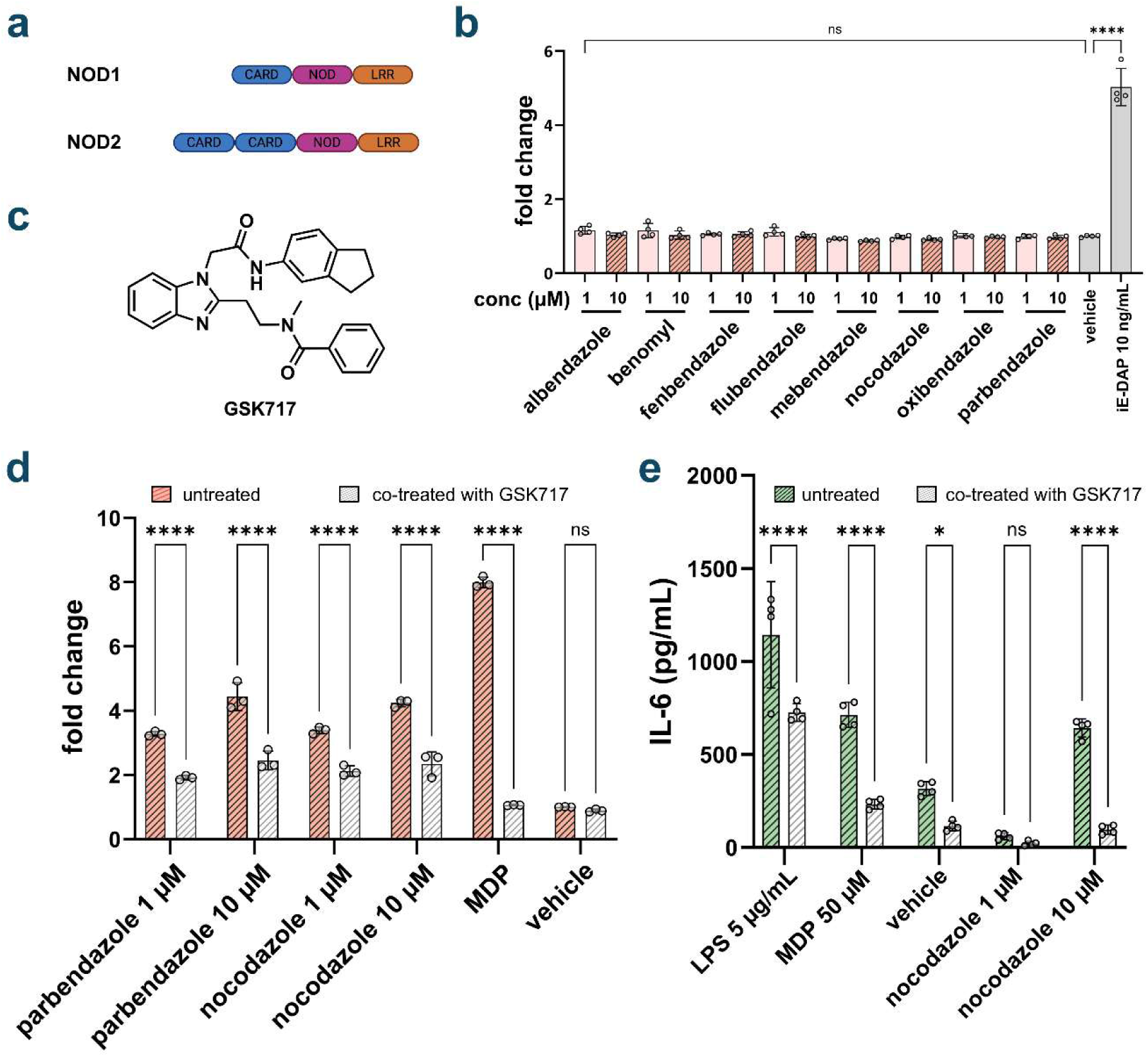
**(a)** Protein domains of NOD1 and NOD2. **(b)** Activation of NOD1 by benzimidazoles. HEK-Blue NOD1 cells were treated with each compound at indicated concentrations for 16 h using a colorimetric assay (n=4). p value determined by one-way ANOVA (ns=not significant, **** p<0.0001). Only one statistical significance bar is shown for clarity. No significance was observed for any compound at either concentration. **(c)** Inhibition of parbendazole and nocodazole by GSK717 in reporter cells. HEK-Blue NOD2 cells were treated with each compound and GSK717 at indicated concentrations for 16 h using a colorimetric assay (n=4). P value determined by one-way ANOVA (ns=not significant, **** p<0.0001). **(d)** Inhibition of cytokine release with GSK717. THP-1 cells were treated with each compound for 20 h and cytokine release was measured by ELISA from cell media. p values determined by one-way ANOVA (ns=not significant, * p< 0.05, **** p<0.0001).

To provide additional confirmation of NOD2-specific signaling, we employed the selective NOD2 inhibitor GSK717 to pharmacologically validate our findings. GSK717 functions as a competitive inhibitor of NOD2 and has been extensively characterized for its ability to inhibit NOD2 activation in both HEK-Blue NOD2 cells and THP-1 cells, exhibiting similar IC50 values across both systems (**Fig 3c**).^12^ Co-treatment with GSK717 resulted in significant inhibition of NOD2 activation induced by the most potent benzimidazole compounds, nocodazole and parbendazole, in HEK-Blue NOD2 cells (**Fig. 3d**). While the degree of inhibition observed with nocodazole was somewhat reduced compared to MDP, the effect remained statistically significant. THP-1 cells, an immortalized human monocytic cell line, provide a physiologically relevant model for assessing NOD2 activation through quantification of cytokine release, particularly IL-8, which serves as a sensitive readout for both NOD2 and NOD1 signaling pathways.^12^ The inhibitory profile observed in HEK-Blue-NOD2 cells was corroborated in THP-1 cells, where GSK717 significantly attenuated nocodazole-induced IL-6 release (**Fig. 3e**). The consistent but incomplete inhibition of benzimidazole-mediated NOD2 activation across both cellular systems suggests that these compounds may engage the NOD2 binding site through a distinct molecular mechanism compared to MDP. Given the competitive nature of GSK717 inhibition, the observed differences in inhibitory efficacy likely reflect unique binding interactions between nocodazole and the NOD2 active site, which may not be fully antagonized by GSK717 under the experimental conditions employed.

### Structural Activity Relationship Studies on Benzimidazoles

Structure-activity relationship (SAR) analysis revealed a critical role for carbonyl functionality in benzimidazole-mediated NOD2 activation. Some of the most potent benzimidazole NOD2 agonists, including mebendazole and nocodazole, share a common structural motif consisting of a carbonyl moiety positioned between two aromatic rings. This structural pattern extends to flubendazole, which also exhibits NOD2 agonistic activity (**Fig. 2a**). In contrast, several benzimidazole analogs bearing sulfone substituents at analogous positions failed to induce NOD2 activation, suggesting that the specific electronic and steric properties of the carbonyl group are essential for biological activity (**Fig. 2b**). To directly assess the functional importance of the carbonyl moiety, we synthesized reduced analogs of flubendazole, mebendazole, and nocodazole wherein the carbonyl groups were converted to their corresponding alcohols (**Fig. 4a**). Evaluation of these reduced compounds in HEK-Blue NOD2 cells demonstrated a dramatic reduction in NOD2 activation, with activity decreased by at least 50% at the highest concentrations tested (**Fig. 4b-d**). More strikingly, at lower concentrations (500 nM and 1 μM), the reduced benzimidazoles were completely inactive, displaying only background-level responses, whereas their parent carbonyl-containing compounds retained measurable NOD2 agonistic activity. These findings establish the carbonyl functionality as a critical pharmacophore element, demonstrating that its presence is essential for dual carbonyl benzimidazole agonists such as mebendazole and nocodazole to activate NOD2.

**Figure 4.**
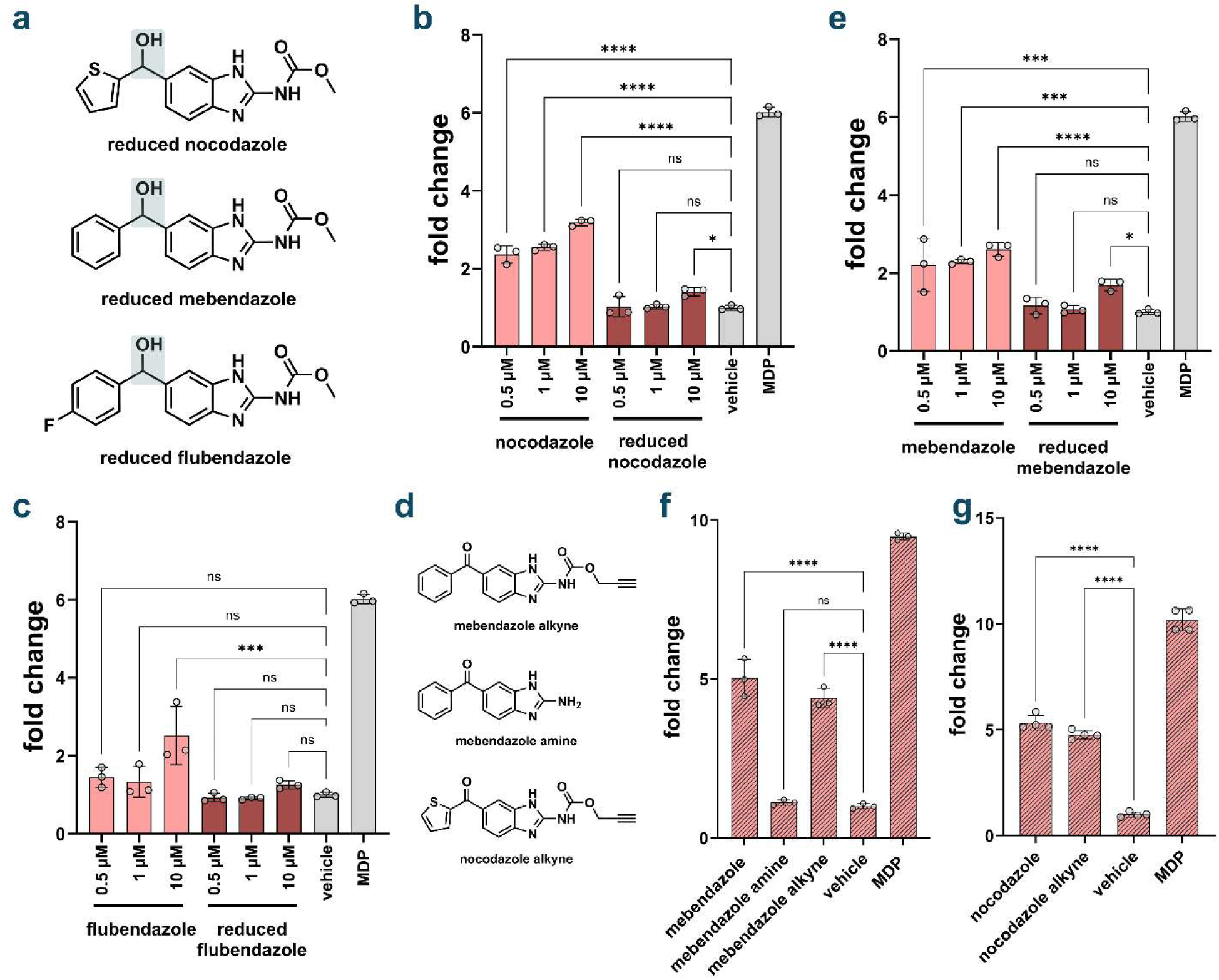
Structural alterations of benzimidazole agonists. **(a)** Chemical structures of reduced benzimidazole compounds and alterations to carbamate group. **(b)** Effect of reduced nocodazole on NOD2 activation. HEK-Blue NOD2 cells were treated with each compound at indicated concentrations 16 h using a colorimetric assay (n=4). p value calculated by one-way ANOVA (ns=not significant, * p<0.1, *** p< 0.001, **** p<0.0001). **C)** Effect of reduced mebendazole on NOD2 activation. HEK-Blue NOD2 cells were treated with each compound at indicated concentrations for 16 h using a colorimetric assay (n=4). **(d)** Effect of reduced flubendazole on NOD2 activation. HEK-Blue NOD2 cells were treated with each compound at indicated concentrations for 16 h using a colorimetric assay (n=4). **(e)** Chemical structures of alterations to carbamate group on nocodazole and mebendazole. **(f)** Impact of carbamate group alteration on mebendazole induced NOD2 activation HEK-Blue NOD2 cells were treated with each compound at 10 mM for 16 h using a colorimetric assay (n=4). **(g)** Impact of carbamate group alteration on nocodazole induced NOD2 activation HEK-Blue NOD2 cells were treated with each compound at 10 mM for 16 h using a colorimetric assay (n=4).

All NOD2 active benzimidazole compounds identified feature a carbamate substituent attached to the benzimidazole core. The only two compounds evaluated that lack this functional group, benomyl and omeprazole, failed to consistently exhibit NOD2 activation (**Fig. 2b**). The carbamate group is metabolically active and could be changed within cells to a different form. To systematically evaluate the functional contribution of the carbamate moiety, we synthesized structural analogs of mebendazole wherein the carbamate was replaced with either an amine or alkyne functionality **(Fig 4e).** The alkyne analog, which preserved the carbonyl component of the original carbamate, maintained comparable NOD2 activation to unmodified mebendazole (**Fig. 4f**). In contrast, the amine analog exhibited dramatically reduced activity, with NOD2 activation diminished to near-background levels. Following these results, and alkyne analog of nocodazole was also synthesized. Nocodazole alkyne also retained similar NOD2 activity to its parent compound, nocodazole, displaying only a slight decrease in signal (**Fig. 4g)**. These structure-activity relationships indicate that while the carbamate group is essential for optimal NOD2 activation by benzimidazoles, the functionality can tolerate modest structural modifications without complete loss of biological activity.

### Benzimidazole Agonists Induce Cytokine Secretion from Immune Cells

The therapeutic potential of NOD2 activation stems primarily from its downstream cytokine signaling cascades. Key inflammatory mediators associated with NOD2 signaling include IL-6 and IL-8, both pro-inflammatory cytokines regulated by NF-κB transcriptional activity.^31,49,50^ We employed THP-1 cells to assess the capacity of benzimidazole compounds to stimulate IL-8 and IL-6 release. THP-1 cells have been validated for quantifying NOD1 and NOD2 activation through measuring cytokine release.^12^ Lipopolysaccharide (LPS) served as an additional positive control, activating Toll-like receptors 2 and 4 (TLR2/TLR4) endogenously expressed in THP-1 cells.^50^ Comparative analysis revealed that nocodazole induced more robust IL-8 secretion than parbendazole (**Fig. 5a**). Interestingly, mebendazole also elicited stronger IL-8 release than parbendazole and maintained activity at lower concentrations more effectively than both nocodazole and parbendazole, despite showing weaker performance in HEK-Blue-NOD2 cells. In THP-1 cells, nocodazole exhibited particularly potent activity, with IL-8 levels frequently approaching the upper detection limit of the ELISA standard curve (**Fig. 5a**). Nocodazole also induced IL-6 secretion, though to a lesser extent than IL-8 (**Fig. 5b**). Reducing the nocodazole concentration to 1 μM led to a marked decrease in both IL-8 and IL-6 secretion, a reduction more pronounced than that observed in HEK-Blue-NOD2 cells (**Fig. 5a-b**). Notably, nocodazole triggered higher cytokine secretion than MDP, even though MDP was used at a 5-fold higher concentration. These results contrast with those from the HEK-Blue-NOD2 reporter assay, in which MDP consistently outperformed the benzimidazole compounds at lower concentrations. The differing NOD2 activation profiles across concentrations and cell types suggest cell-type-specific responses to benzimidazoles and MDP, or differences in downstream biochemical processes.

**Figure 5.**
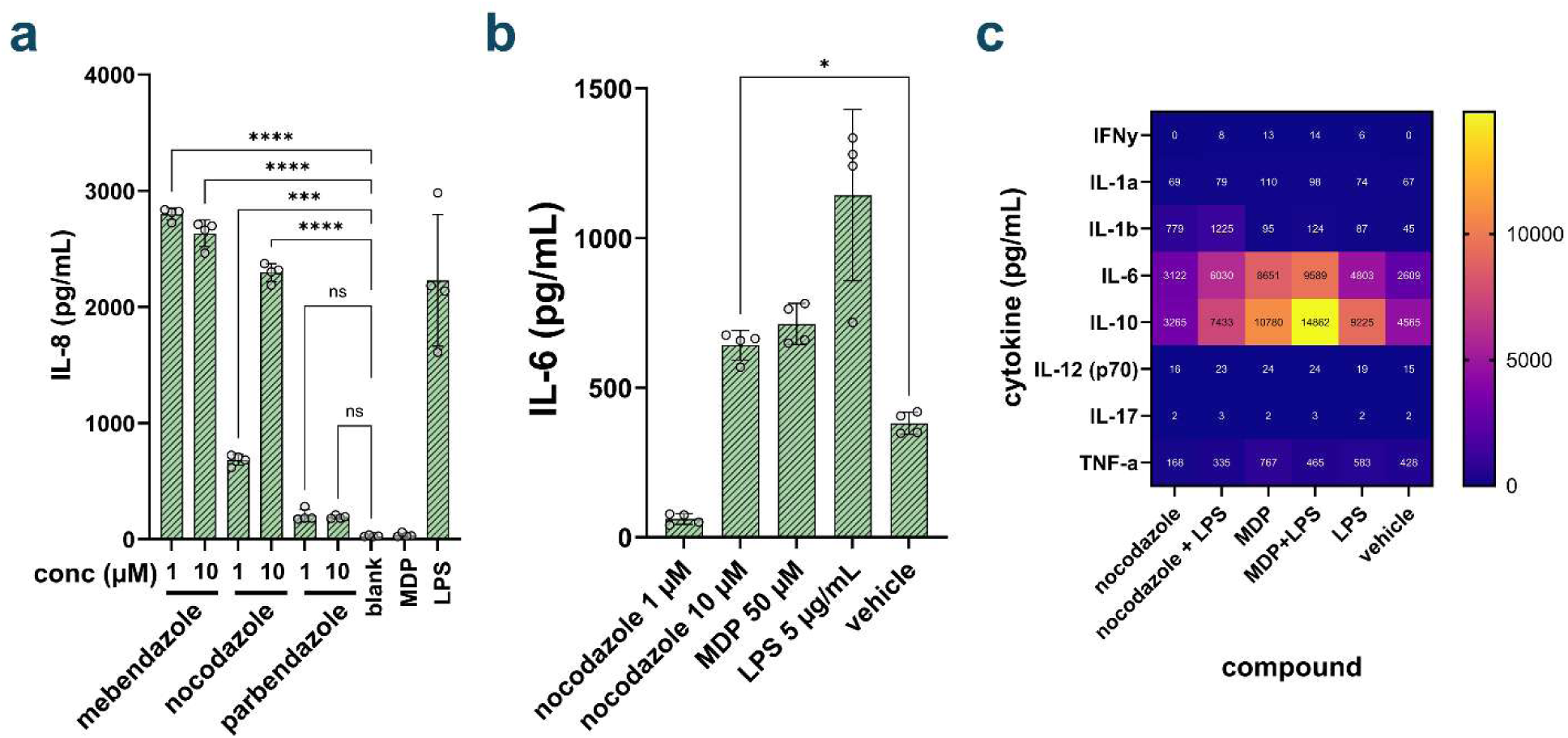
**(a)** Release of IL-8. THP-1 cells were treated with each compound for 20 h and cytokine release was measured by ELISA from cell media (n=4). p value calculated by one-way ANOVA (ns=not significant, *** p<0.001 **** p<0.0001). **(b)** Release of IL-6. THP-1 cells were treated with each compound for 20 h and cytokine release was measured by ELISA from cell media (n=4). P value calculated by one-way ANOVA (* p<0.1). **(c)** Luminex assay results. Mouse BMDMs were treated with each compound overnight at 20 µM, with 10 ng/mL LPS when applicable. Cytokine levels were measured by Luminex.

Beyond individual cytokine quantification, we sought to determine whether benzimidazole agonists generate the broader pro-inflammatory cytokine signature characteristic of MDP-induced NOD2 activation. Extensive profiling is critical, as the multifaceted therapeutic effects of NOD2 activation depend on the comprehensive immune response to multiple cytokines released rather than a single mediator.^1,4,5,26^ We therefore employed LUMINEX multiplex immunoassays to analyze supernatants from mouse bone marrow-derived macrophages (BMDMs) stimulated with benzimidazoles or MDP with LPS. The results demonstrate that MDP and benzimidazole agonists produce remarkably similar cytokine release patterns (**Fig. 5c**). Both compound classes strongly induced IL-6 and IL-10 secretion, cytokines canonically associated with NOD2 signaling.^1,33^ Notably, nocodazole elicited more robust IL-1β release compared to MDP. Given that IL-1β represents another critical mediator of NOD2-dependent responses, this enhanced induction suggests potentially superior therapeutic efficacy. Collectively, these data indicate that nocodazole can engender the complex cytokine milieu associated with physiological NOD2 activation.

## CONCLUSION

We have identified and characterized a novel class of FDA-approved benzimidazole compounds as potent NOD2 agonists. Among them, nocodazole, demonstrating nanomolar potency, emerged as the lead compound. Pharmacological validation confirmed that nocodazole activates NOD2-specific signaling and induces a cytokine release profile consistent with that triggered by the natural ligand MDP. Importantly, nocodazole features a scaffold structurally distinct from peptidoglycan-derived NOD2 agonists, offering the potential for improved pharmacological properties and reduced side effects. Given NOD2’s established roles in enhancing cancer immunotherapy and mitigating inflammatory bowel disease, these benzimidazole-based agonists represent promising therapeutic candidates. The discovery of nocodazole and related compounds also provides valuable tools for mechanistic studies of NOD2 biology and for the development of synthetic small-molecule immunomodulators.

## ACKNOWLEDGEMENT

This study was supported by the NIH grant 1R01AI178975-01 (M.M.P.), R35GM124893 (M.M.P.), R01AI179080-01 (M.M.P.).

