## Supplementary Information for "Identification and Evaluation of Benzimidazole-Agonists of Innate Immune Receptor NOD2"

Figure S1........................................................................**Error! Bookmark not defined.**
Figure S2........................................................................**Error! Bookmark not defined.**
Figure S3........................................................................**Error! Bookmark not defined.**

#### SUPPORTING FIGURES

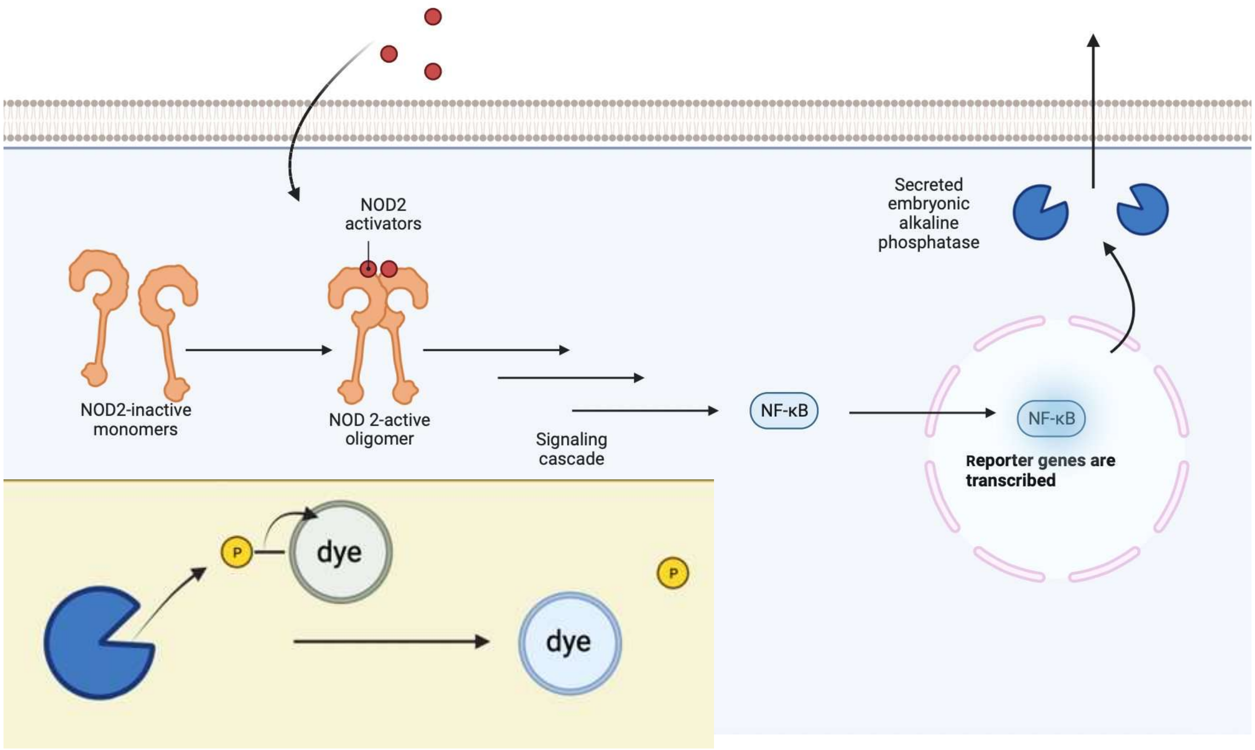

**Fig. S1 Function of NOD2-HEK-Blue reporter cells.** NOD2 activators cause the release of SEAP into the media, leading to a color change as a phosphate is removed from the dye.

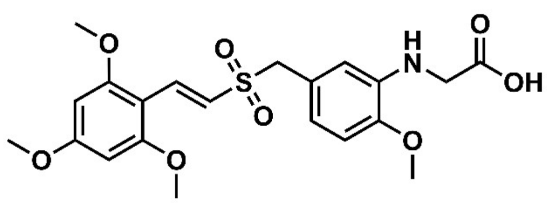

**rigosertib**

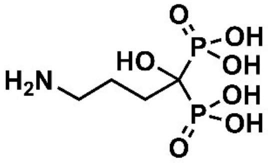

**alendronate**

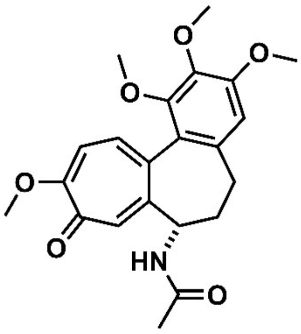

**colchicine**

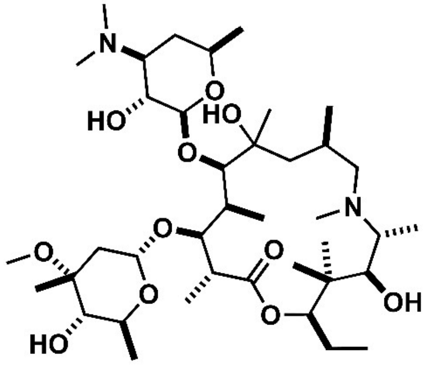

**azithromycin**

**Fig. S2 Chemical structures of other screen hits.** Chemical structures of rigosertib, alendronate sodium, colchicine, and azithromycin.

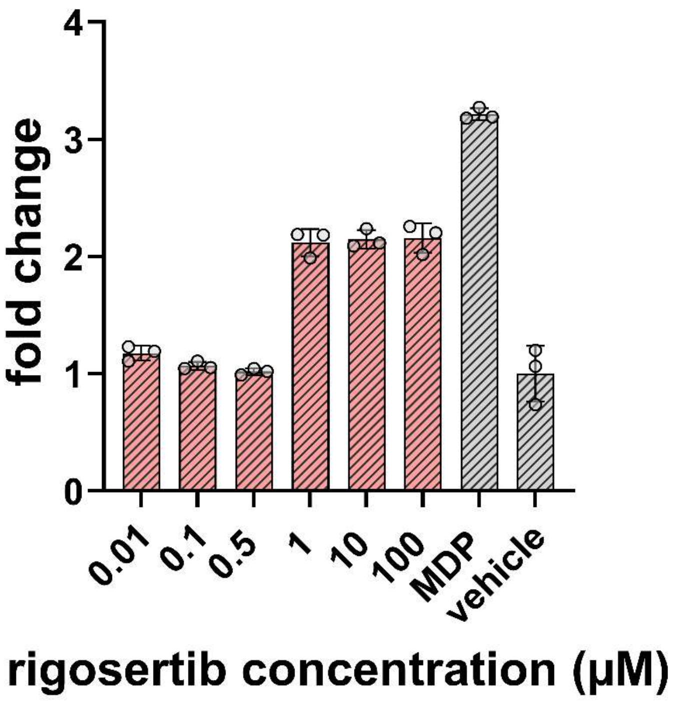

**Fig. S3 Dose dependance of rigosertib in HEK-Blue NOD2 cells**. HEK-Blue cells were treated with rigosertib in DMSO for 16 h using a colorimetric assay. The analysis was performed by measuring the absorbance of each well at 655 nm. Fold change calculated by compound absorbance/average background (n=3).

### General Experimental Details

##### Cell Culture

HEK-Blue-NOD2 cells (InvivoGen, hkb-hnod2v2) were grown in DMEM (DMEM, Sigma Aldrich, D6429) supplemented with 10% fetal bovine serum (FBS) (FBS, Thermo Scientific, A5670801) and 1% penicillin/streptomycin (P/S, Sigma Aldrich, P4333). THP-1 (ATCC, TIB-202) cells were grown in RPMI-1640 media (RPMI-1640, Fisher Scientific, 11-875-119) supplemented with 10% fetal bovine serum (FBS) and 1% penicillin/streptomycin. Bone marrow derived macrophages (BMDM) were obtained from the Bullock Lab. Cells were incubated at 37 $^oC$ with 5% $CO_2$.

##### Screen of the L10121 Discovery Probe FDA Approved Library

When HEK-Blue cells reached 80-90% confluency, they were removed from their flasks with Trypsin/EDTA (ThermoFisher, 25200056) and resuspended in HEK-Blue detection media (HEK-Blue Detection, InvivoGen, hb-det2). Cells were then plated at 50,000 cells/well on a 96-well plate. Test compounds library plates were thawed 30 minutes beforehand, and DMSO solutions were pipetted directly into assay plates and then mixed with cells via pipetting. Cells were incubated at 37 $^oC$ and 5% $CO_2$ for 16 hours, and subjected to measure absorbance at 655nm on a UV-Vis plate reader. 1 μM MDP (N-Acetylmuramyl-L-alanyl-D-isoglutamine hydrate, Sigma Aldrich, A9519) and 2 μL of DMSO were used as positive and negative controls, respectively.

##### HEK-Blue NOD2 Assay

When HEK-Blue cells reached 80-90% confluency, they were removed from their flasks with Trypsin/EDTA and resuspended in HEK-Blue detection media. Benzimidazole compounds (see materials) were diluted sequentially from DMSO stocks into acetonitrile and then into HEK-Blue Detection media to obtain desired final concentration. Cells were then plated at 50,000 cells/well on a 96-well plate and treated with compounds for 16hrs, incubating at 37 $^oC$ and 5% $CO_2$. Data was acquired at 655nm on a UV-Vis plate reader.

**ELISA**

When THP-1 cells reached a confluence in the flask approaching 8 x $10^6$ cells/mL, they were spun down at 1000 x g for 5 min, resuspended in fresh growth media, and plated at 500,000 cells/well on a 24-well plate. Benzimidazole compounds were diluted sequentially from DMSO stocks into acetonitrile and then into RPMI media. Cells were incubated with compounds for 20-24hrs. The cell media was extracted and used for IL-8 and IL-6 ELISAs (ELISA-MAX Deluxe Set Human IL-8/IL-6, BioLegend, 431504/430504). The ELISA was conducted following the procedure laid out in the product information for the BioLegend Deluxe ELISA kit.

**BMDM isolation**

Mouse BMDMs were derived from BM isolated from the femurs and tibias of 8- to 10-week-old C57BL/6J mice and cultured in RFHP10 media (RPMI 1640 supplemented with 10% heat-inactivated FBS, 10 mM *N*-2-hydroxyethylpiperazine-*N′*-2-ethanesulfonic acid, and 1% penicillin–streptomycin–L-glutamine) supplemented with 20 ng/mL of M-CSF for 10 days. Media was replaced with fresh RFHP10 supplemented with 20 ng/mL of M-CSF on day 5 and BMDMs were used from day 6-10.

**LUMINEX**

BMDMs were removed from their growth plates with Trypsin/EDTA and resuspended in RFHP10 supplemented with M-CSF. Compounds were added at a concentration of 20 μM. Benzimidazole compounds were diluted sequentially from DMSO stocks into acetonitrile and then into DMEM media. A fresh stock of LPS from E. coli was added at 10 ng/mL in relevant conditions. BMDMs incubated at 37 $^{o}$C and 5% $CO_2$ overnight with the compounds and controls. Media was removed and frozen at –80 $^{o}$C for storage. The media was thawed and sent off for LUMINEX at the UVA flow core.

**Inhibition by GSK717**

HEK-Blue NOD2 cells were grown in appropriate media (listed above). The cells were grown to approximately 80% confluency, removed from their flasks with Trypsin/EDTA and resuspended in HEK-Blue detection media. Benzimidazole compounds were diluted sequentially from DMSO stocks into acetonitrile and then into HEK-Blue Detection media. GSK717 (MedChem Express, HY136555) was diluted from DMSO stock into ethanol and then added to the plate directly at the same time as compounds and positive controls. Cells were then plated at 50,000 cells/well on a 96-well plate and treated with compounds for 16hrs, incubating at 37 $^{o}$C and 5% $CO_2$. Results were read at 655nm on a UV-Vis plate reader.

#### Thin Layer Chromatography

Thin layer chromatography analyses for monitoring reaction progress were performed on aluminum backed thin layer silica gel plates (Merck F254), plates were developed in appropriate solvents mentioned in individual reaction and were observed under UV lamp. Column chromatography on silica gel (Supelco, 60 Å, 230-400 mesh, 40-63 m particle size) was performed on manually packed column with selected solvents for each individual sample.

#### HPLC

Preparative reverse phase HPLC purification was performed on instruments equipped with Waters 1525 pumps and 2489 UV/Visible Detector on a Phenomenex Luna 10 μm C8(2) 100 Å (250 x 21.2 mm) or C18 columns using a 5 to 100% linear gradient of methanol in H2O or MeCN in water each containing 0.1% TFA at 10 mL/min. The HPLC fractions of the desired compounds were first concentrated under reduced pressure using a rotary evaporator. The concentrated aqueous solutions were lyophilized with Labconco Freezone 4.5L lyophilizer (-84 $^{o}$C). The purity of the samples was ascertained by either TLC or analytical HPLC using a Phenomenex Luna 5 μm C8(2) 100 Å (250 x 4.6 mm) on the same instrument; using gradient elution in $H_2O$/CH3CN or H2O/MeOH with 0.01% TFA in each solvent at 1 mL/min prior to its use in biological assays. Some of the compounds having amino function isolated from HPLC may have been trifluoroacetate salt form and these were not neutralized prior to characterization by methods described below and were used as is for bioassays.

#### NMR

$^{1}H$ and $^{13}C$-NMR spectra for final compounds and intermediates were acquired on a Varian 600MHz spectrophotometer. All NMR spectra were processed and analyzed using MestreNova software. Deuterated solvents were used as received from Cambridge Isotopes. Residual solvent signal from $CDCl_3$, $CD_3OD$ and DMSO-$d_6$ referenced to tetramethylsilane (TMS) were used as reference standards for defining chemical shifts. Chemical shifts are reported in $\delta$ ppm and coupling constants ($J$) are reported in Hertz [Hz]. Mass analysis for follow up of reaction or final product analysis was performed on Advion Expression® CMS mass spectrometer using standard ESI parameters for intermediates and final products. For the analysis of fragmentation sensitive compounds, low fragmentation, low energy setup was used.

**Mass Spectrometry**

MALDI-TOF mass spectra were obtained for certain high molecular weight compounds on the Shimadzu MALDI-8020 instrument with α-Cyano-4-hydroxycinnamic acid (α-CHCA) matrix. The observed molecular weights for compounds were represented as m/z. Additional LCMS analyses were performed on Shimadzu Prominence-*i* LCMS system equipped with LC-2030C 3D liquid chromatograph, autosampler, PDA and LCMS-2020 mass detector. Data analysis was performed with Lab Solutions software. High resolution electrospray ionization mass spectrometry (HRMS, ESI/MS) analyses were obtained in positive ion mode on an Agilent 6545B Q-TOF LC/MS equipped with 1260 Infinity II LC system with auto sampler and equipped with C18(2) column (Luna 5 µm 100Å 250 x 4.6 mm). The LCMS/MS data was processed with Mass Hunter software, and subsequent quantification was done with Q-Tof MS quantitative software. UV spectroscopic analysis was performed on Genesys 50 (Thermo Scientific) UV-Visible Spectrophotometer.

**Statistical Analysis**

Unless otherwise specified, statistical analysis was conducted using GraphPad Prism 9.5. One-way ANOVA was used to calculate statistical significance. Bars represent standard deviation.

### Materials

**General information about materials and instruments:**

Chemicals, reagents, and solvents used for the synthesis were purchased from standard sources such a Sigma-Aldrich (St. Louis, MO, USA), Fisher Scientific (Hampton, NH, USA), VWR (Randor, PA, USA) and Alfa Aesar (Ward Hill, MA, USA). Chemicals and reagents were used as is after acquisition, purity of compounds for biological testing were assigned to be >95%. The stock solutions of these compounds were made in DMSO at 10-20 mM and stored at -20 $^{o}$C.

**Materials**

N-acetylmuramulyl-L-alanyl-D-isoglutamine hydrate (MDP): Sigma-aldrich A9519

Oxibendazole: Ak Scientific E369

Rigosertib: Ak scientific 4351EQ
Colchicine: AK scientific J10109

GS9973: AK scientific 2440AH

GSK717: Medchem Express HY-136555

Mebendazole: AK scientific E711

Albendazole: Cayman Chemicals 23705

Ricobendazole: Cayman Chemicals 21880

Flubendazole: Cayman Chemicals 26064

Nocodazole: Cayman Chemicals 13857

Carbendazim: Cayman Chemicals 23852

Parbendazole: Medchem Express HY115364

Albendazole Sulfone: Cayman Chemicals 35445

Fenbendazole Sulfone: Cayman chemicals 20921

Oxfendazole: Cayman Chemicals 29742

Benomyl: Cayman Chemicals 34634

#### Chemical Synthesis

##### Hydrolysis of 5-Aroyl-2-benzimidazolecarbamic acid methyl ester to 2-Amino-5-aroyl-1*H*-benzimidazole:

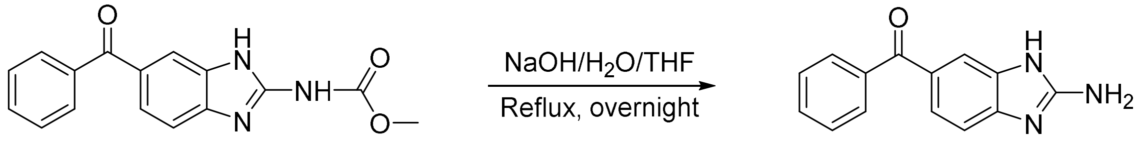

Representative example: Methyl(6-benzoylbenzo[d]thiazol-2-yl) carbamate (Mebendazole, 312 mg, 1.05 mmol) suspended in 5% 1M, NaOH and THF (1:9, 25mL). The heterogenous reaction mixture heated on oil to reflux conditions overnight (~15 hrs). The next day, tlc analysis indicated starting material consumed from a new polar compound. The reaction mixture concentrated under reduced pressure using a rotary evaporator. The left-over residue was carefully acidified with 0.1M HCl to pH 8.0 and extracted with ethyl acetate (10mL x3), The combined organic layer was washed with saturated. $NaHCO_3$ and finally with brine, dried over $Na_2SO_4$ and concentrated to yield off-white solid (212 mg, 85%), 1H-NMR matched the reported data;[1] purity of sample was confirmed with HPLC.

Mebendazole-amine $^1$H-NMR (DMSO-$d_6$) δ 7.62 (m, 8H), 8.86 (s, 2H), 12.89 (bs, 2H); $^{13}$CNMR (75 MHz, DMSO-$d_6$) δ 111.1, 113.1, 125.6, 128.5, 129.4, 129.8, 131.7, 132.3, 133.3,137.5,151.6, 194.8; ESI-MS: calculated *m/z* [M+H] + 238.09, observed *m/z* 238.1. Fenbendazole-amine $^1$H-NMR (DMSO-$d_6$) δ = 7.71-7.64 (m, 2H), 7.64-7.48 (m, 4H), 7.40 (d, *J* = 1.7 Hz, 1H), 7.20 (d, *J* = 8.2 Hz, 1H), 6.74 (s, 2H). ESI-MS: calculated *m/z* [M+H] + 242.31, observed *m/z* 242.31

Nocadozole-amine ESI-MS: calculated *m/z* [M+H] + 243.09, observed *m/z* 243.1.

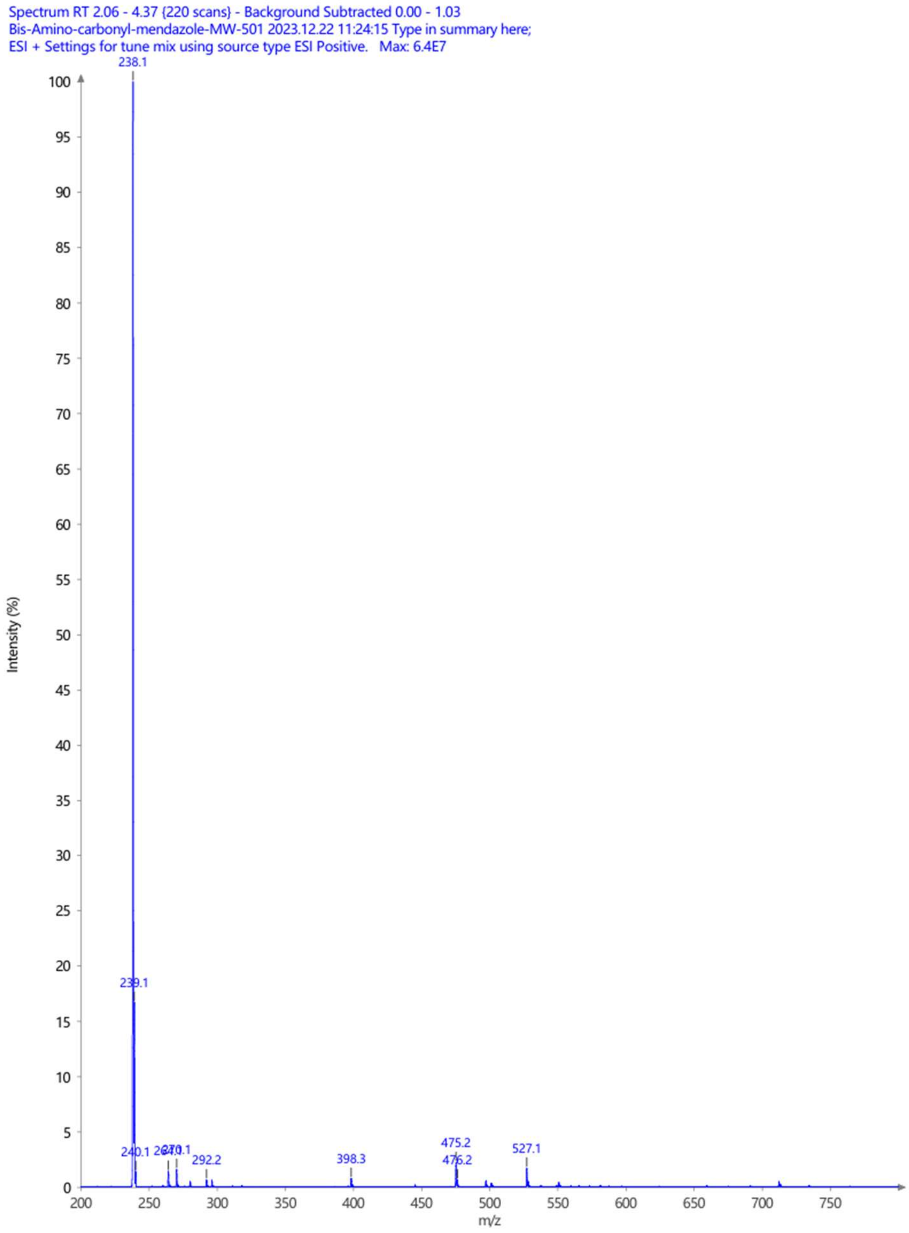

Mass spectrum of mebendazole derived amine, m/z 238 observed for $[M+H]^+$

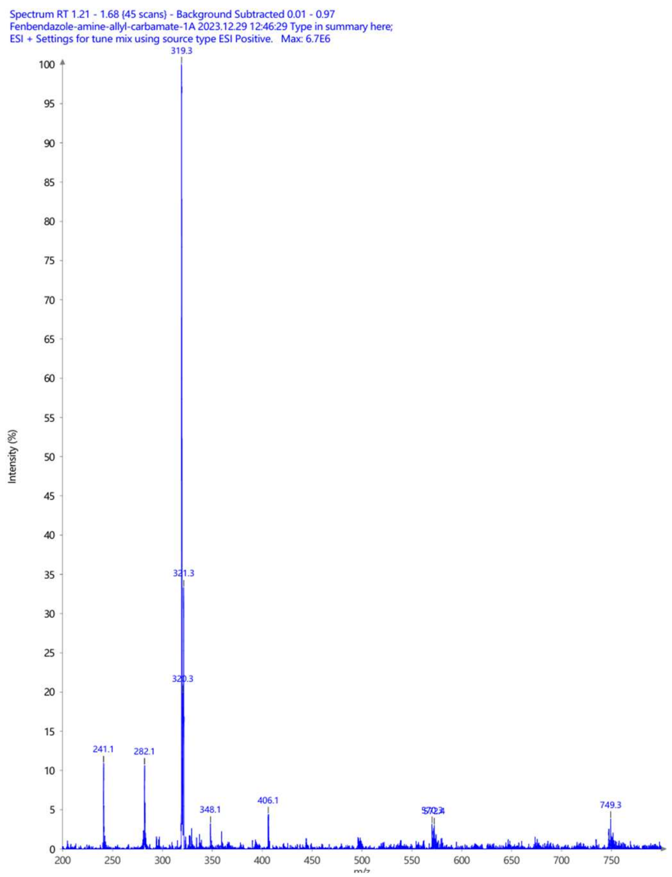

Mass spectrum of Fenbendazole derived amine, m/z 319 observed for $[M+H]^+$

**Reduction of (5-aroyl-1*H*-benzimidazol-2-yl)carbamic acid methyl ester derivatives with $NaBH_4$:**

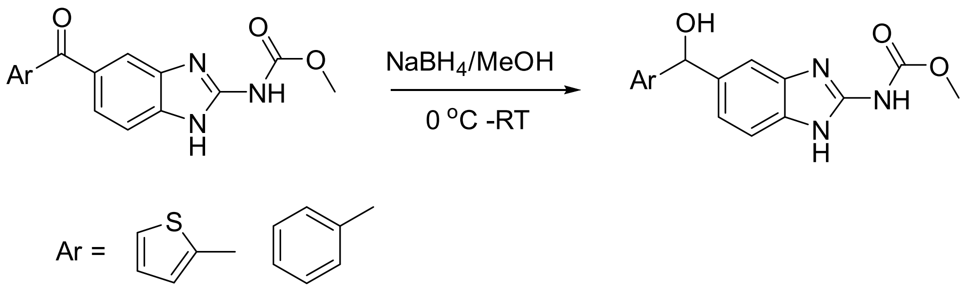

The reduction was carried out using modified reported conditions[2] as follows:

The starting aroyl-benzimidazole compounds (2 mmol) were dissolved in 20 mL of methanol and cooled in ice water to this solution were added $NaBH_4$ (80.0 mg, 2 mmol, 4 eq.) while stirring. The reaction mixture was allowed to warm to RT over 30 minutes and further stirred for an additional 30 min. TLC analysis indicated all starting material consumed to form polar product ($CHCl_3$:MeOH, 98:2). The volatiles were removed under reduced pressure using rotary evaporator and the leftover residue was washed with ether and acidified gently with dil. HCl to ensue precipitation. Water (5 mL) was added and the suspension transferred to a plastic tube for centrifugation. The white solid settled forms a pellet; the aqueous supernatant was removed. The solid pelleted residue was resuspended in DI water and centrifuged again. The process was repeated two more times to remove all water-soluble byproducts and impurities. Finally, the left over solid was frozen in -80 $^{o}$C and lyophilized to yield solid. $^1$H-NMR and mass spectroscopy data matched literature reported data confirming the formation of desired alcohols (Yields: 60-75%) (ref).

**Synthesis of [5-(2-thionyl)-1*H*-benzimidazol-2-yl)carbamic acid prop-2-yn-1yl ester or Prop-2-yn-1-yl(5-(thiophene-2-carbonyl)-1*H*-benzo[*d*]imidazole-2yl)carbamate**:

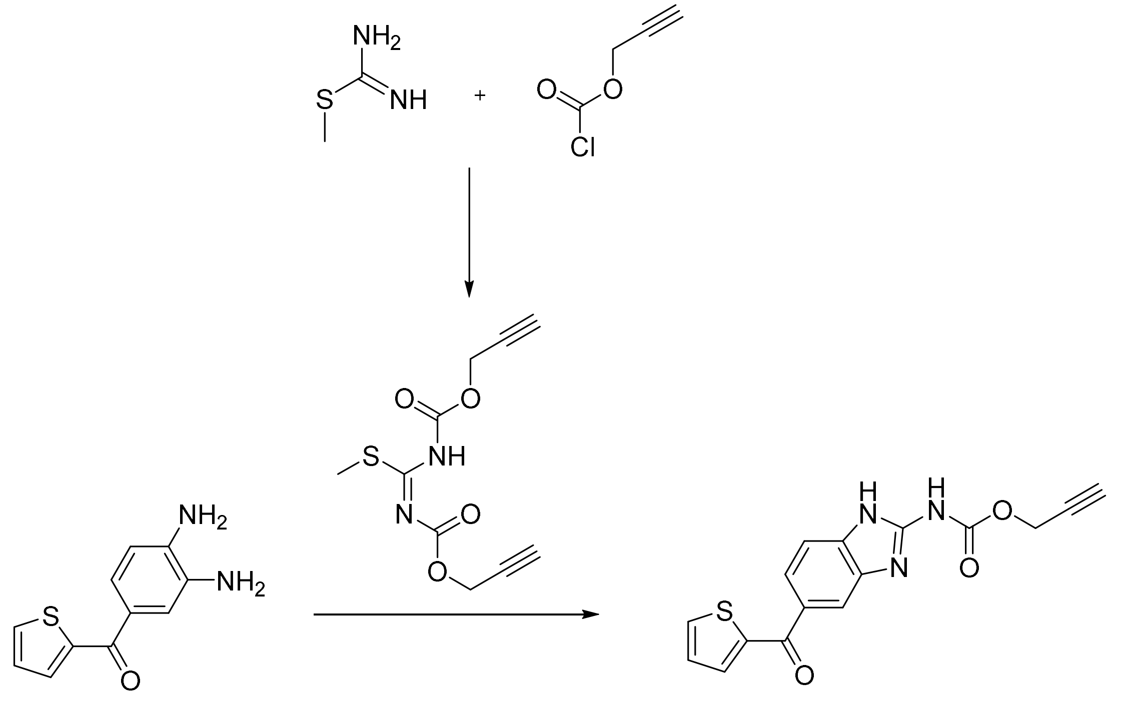

Following the reported procedure for synthesis of bendazole-carbamate synthesis,[3] we used propargyl-chloroformate to make the first 2-propyn-1-yl *N*-[(methylthio)[[(2-propyn-1-yloxy)carbonyl]amino]methylene]carbamate.

S-methylthiuronium sulfate (5.00 g, 35.9 mmol, 1.00 equiv.) suspended in $CH_2Cl_2$ (50 mL), and aqueous $NaHCO_3$ (10%) solution (150 mL) and cooled in ice water bath (5 oC). To this was added solution of propargyl chloroformate (11.5 mL, 108 mmol, 3.00 equiv. in $CH_2Cl_2$, 20 mL) dropwise and stir the mixture at room temperature for 3 h. The organic $CH_2Cl_2$ layer separated, and aqueous layer was extracted once $CH_2Cl_2$. The combined $CH_2Cl_2$ layers washed with water, brine and dried over $Na_2SO_4$. Removal of volatiles using rotary evaporator under reduced pressure afforded transparent syrup, which upon trituration with ether gave solid. White solid (5.8g, 63%), $^1$H-NMR, Mass spect: $^1$H-NMR ($CDCl_3$) $\delta$ 2.45 (s, 3H, SMe), 2.55 (t, *J* =1Hz, 1H), 2.51 (t, *J* =1Hz, 1H), 4.78 (d, *J* =1Hz, 2H), 4.75 (d, *J* =1Hz, 2H), 11.83 (brs, 1H, NH). ESI-MS: calculated *m/z* $[M+H]^+$ 255.26, observed *m/z* 255.1 and [M-15] for methyl loss m/z 239.0

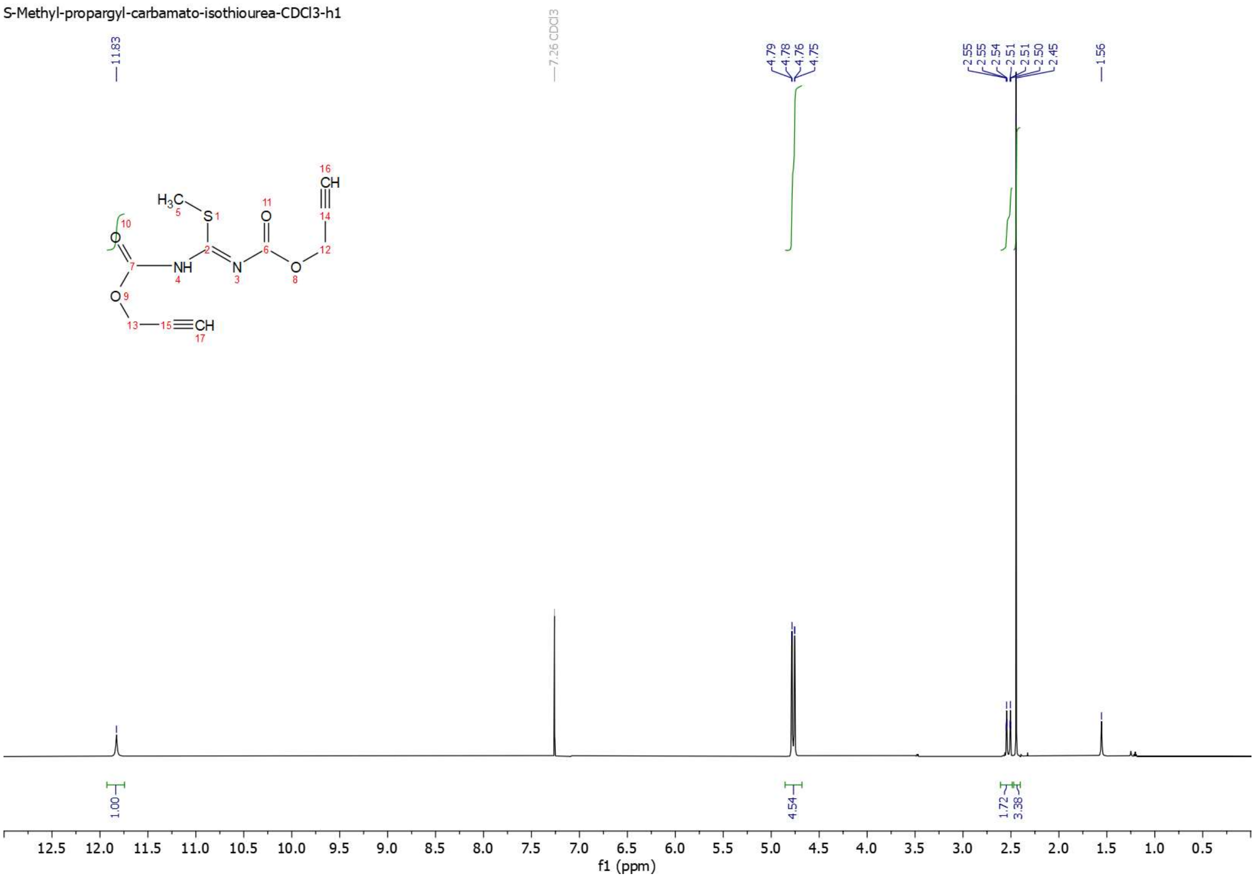

**Figure:** [1]H-NMR of 2-Propyn-1-yl N-[(methylthio)[[(2-propyn-1-yloxy)carbonyl]amino] methylene] carbamate.

**Synthesis of propargyl-carbamate derivatives of bendazole:**

Example- Prop-2-yn-1-yl(6-(thiophene-2-carbonyl)-1H-benzo[d]imidazol-2-yl)carbamate A suspension of (3,4-Diaminophenyl)-2-thienylmethanone (220 mg, 1.0 mmol) and 2-propyn-1-yl*N*-[(methylthio)-[(2-propyn-1-yloxy)carbonyl]amino]methylene]carbamate (275.0mg, 1.08 mmol) in acetic acid:water (3:7, 10 mL) was heated at 100 $^{o}$C for 5 hr in an oil bath. The reaction mixture was then allowed to cool to room temperature and basified with saturated $NaHCO_3$ then extracted with chloroform (3 x 15 mL). The organic layer was combined washed with brine, dried over anhydrous $Na_2SO_4$ subsequent concentration under reduced pressure using rotary evaporator afforded solid (247 mg, 76%). NMR sample contains water. $^{1}$H-NMR (DMSO-$d_6$) δ 3.59 (t, *J*=1.0Hz, 1H, CH), 4.86 (d, *J* =1Hz, 2H, $CH_2$), 7.29 (dd, *J*= 1 and 6Hz, 1H, ArH), 7.53 (d, J= 6Hz, 1H, ArH), 7.66 (dd, *J*= 1 and 6Hz, 1H, ArH), 7.75 (d, *J*=1 Hz, 1H, ArH), 7.96 (s, 1H, ArH), 8.05 (d, J= 6

Hz, 1H, ArH). Partial $^{13}$C-NMR (DMSO-$d_6$) δ 53.0, 78.0, 78.6, 123.1, 128.6, 134.7, 134.7, 143.4, 186.9. ESI-MS: calculated *m/z* $[M+H]^+$ 326.0, observed *m/z* 326.0.

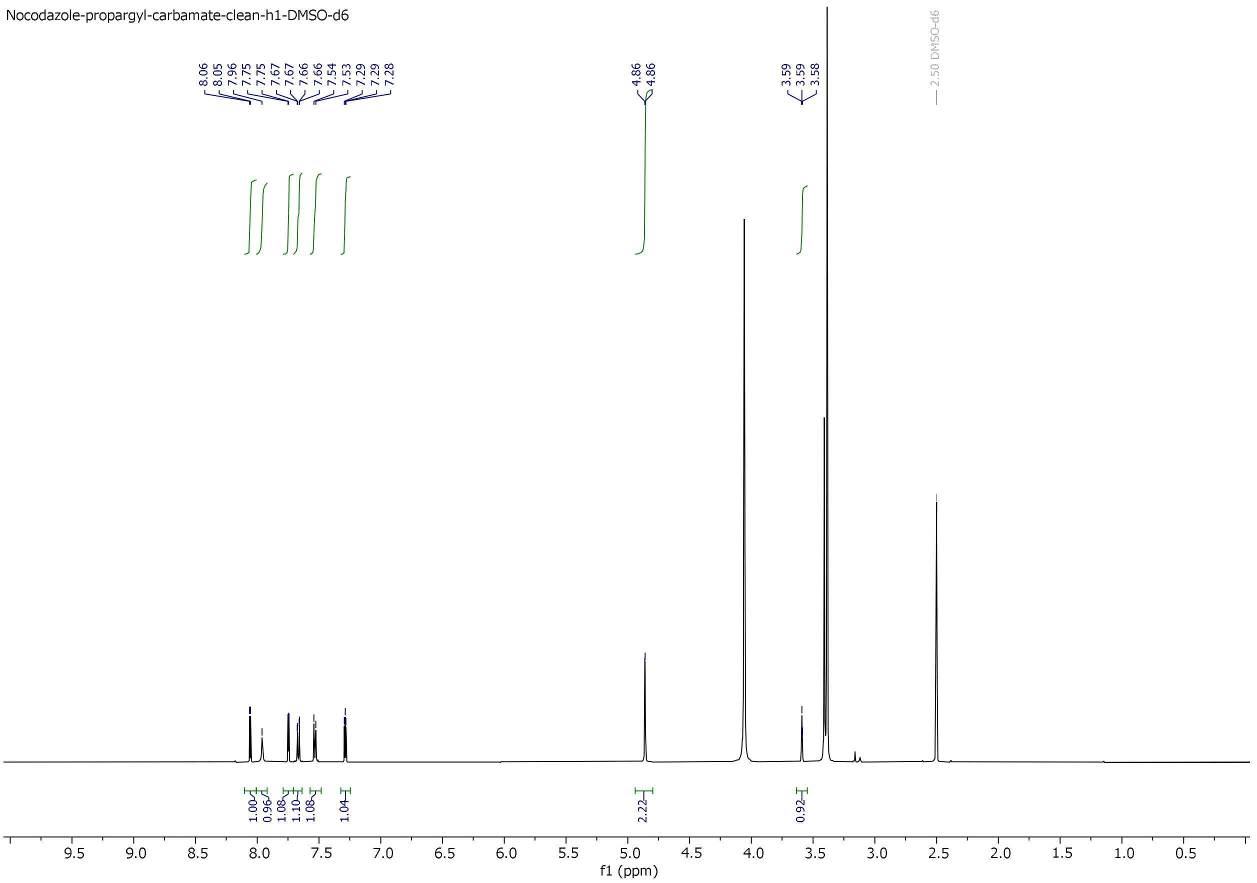

**Figure**: $^1$H-NMR of Prop-2-yn-1-yl (6-(thiophene-2-carbonyl)-1H-benzo[d]imidazol-2-yl) carbamate

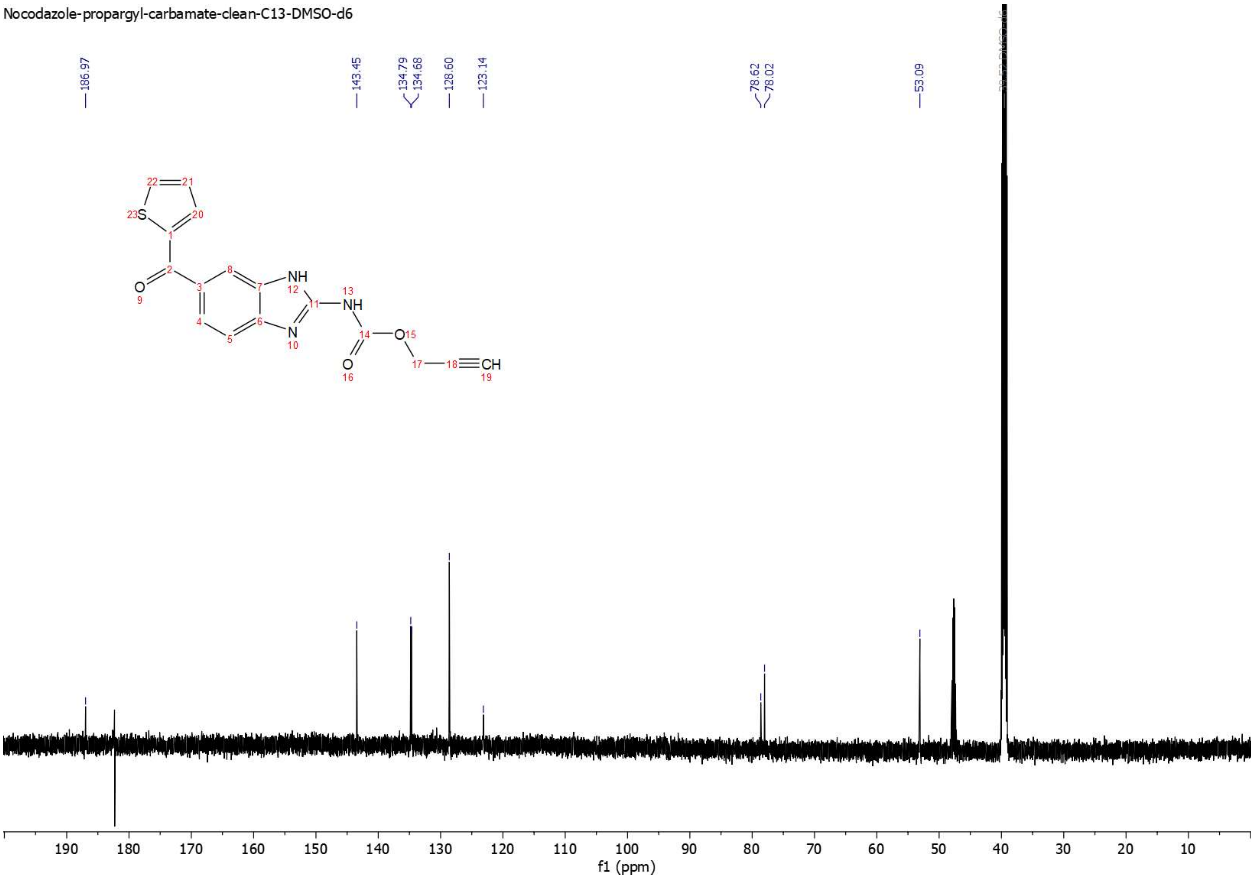

**Figure**: $^{13}$C-NMR of Prop-2-yn-1-yl (6-(thiophene-2-carbonyl)-1H-benzo[d]imidazol-2-yl) carbamate

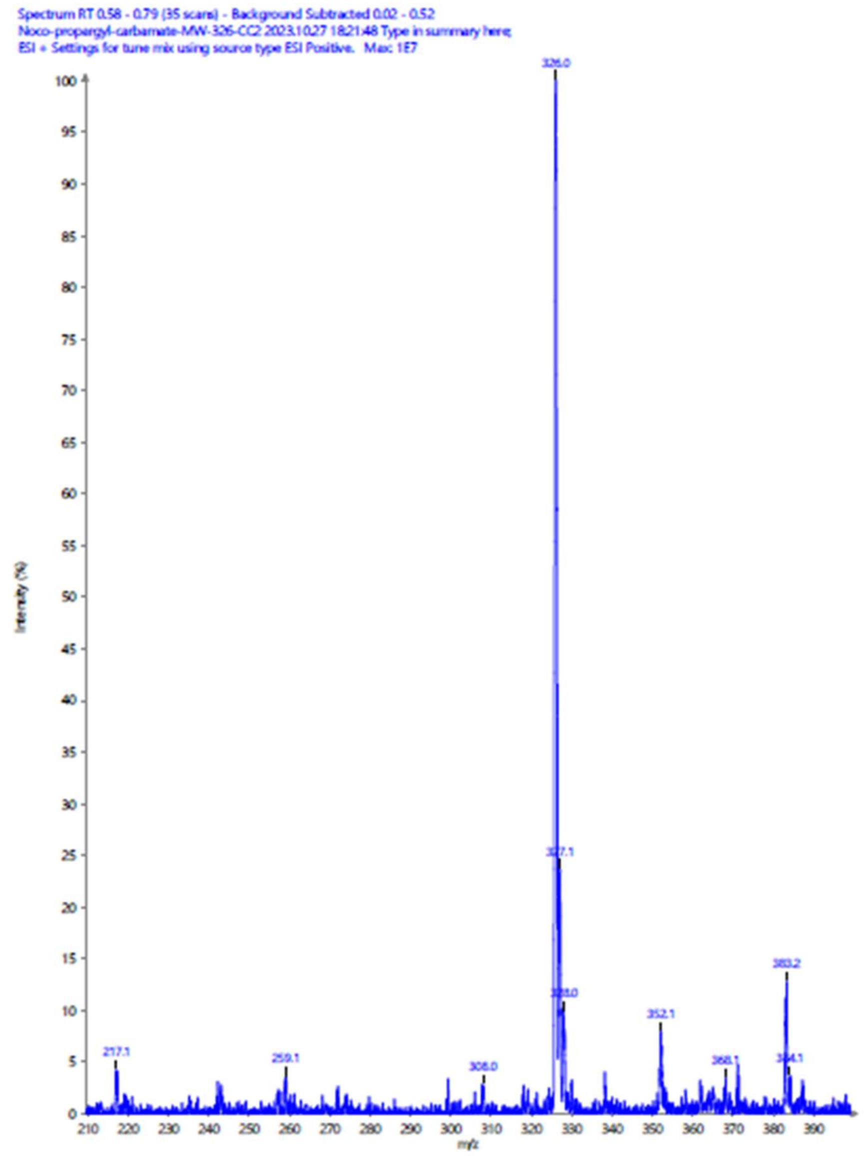

Characterization data for Prop-2-yn-1-yl (6-(benzene-2-carbonyl)-1H-benzo[d]imidazol-2-yl) carbamate:

$^{1}$H-NMR (DMSO-$d_6$) δ 3.62 (t, *J*=1.0Hz, 1H, CH), 4.86 (d, *J* =1Hz, 2H, $CH_2$), 7.52-7.64 (m, 4H, ArH), 7.68 (t, *J*=6 Hz, 1H, ArH), 7.71 (d, *J*= 6Hz, 2H, ArH), 7.85 (s, 1H, ArH), 12.04 (brs, 1H, NH). Partial $^{13}$C-NMR (DMSO-$d_6$) δ 53.0, 78.0, 78.5, 123.8, 128.3, 129.1, 131.8, 138.3, 143.4, 195.5. ESI-MS: calculated *m/z* $[M+H]^+$ 320.3., observed *m/z* 320.2.

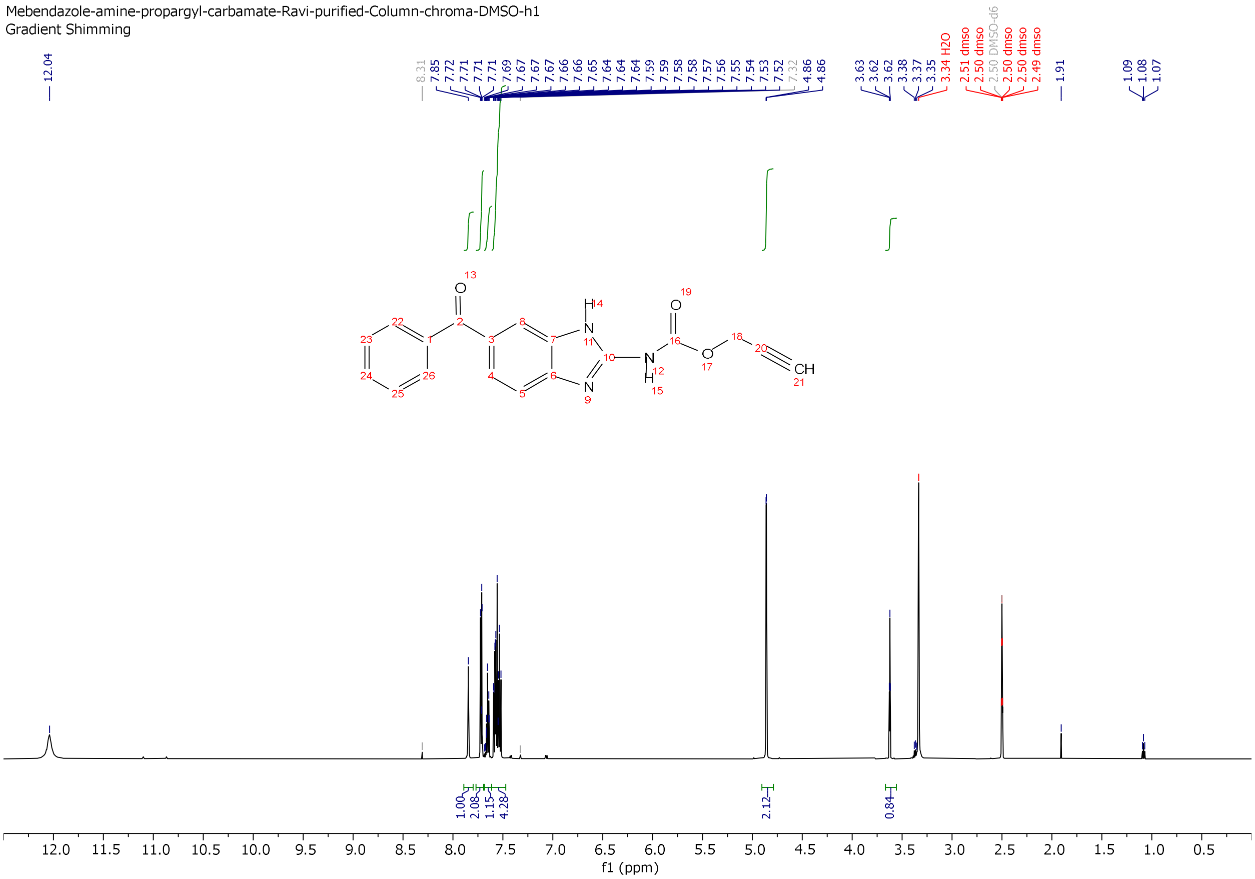

**Figure**: $^{1}$H-NMR of Prop-2-yn-1-yl (6-(benzene-2-carbonyl)-1H-benzo[d]imidazol-2-yl) carbamate

?

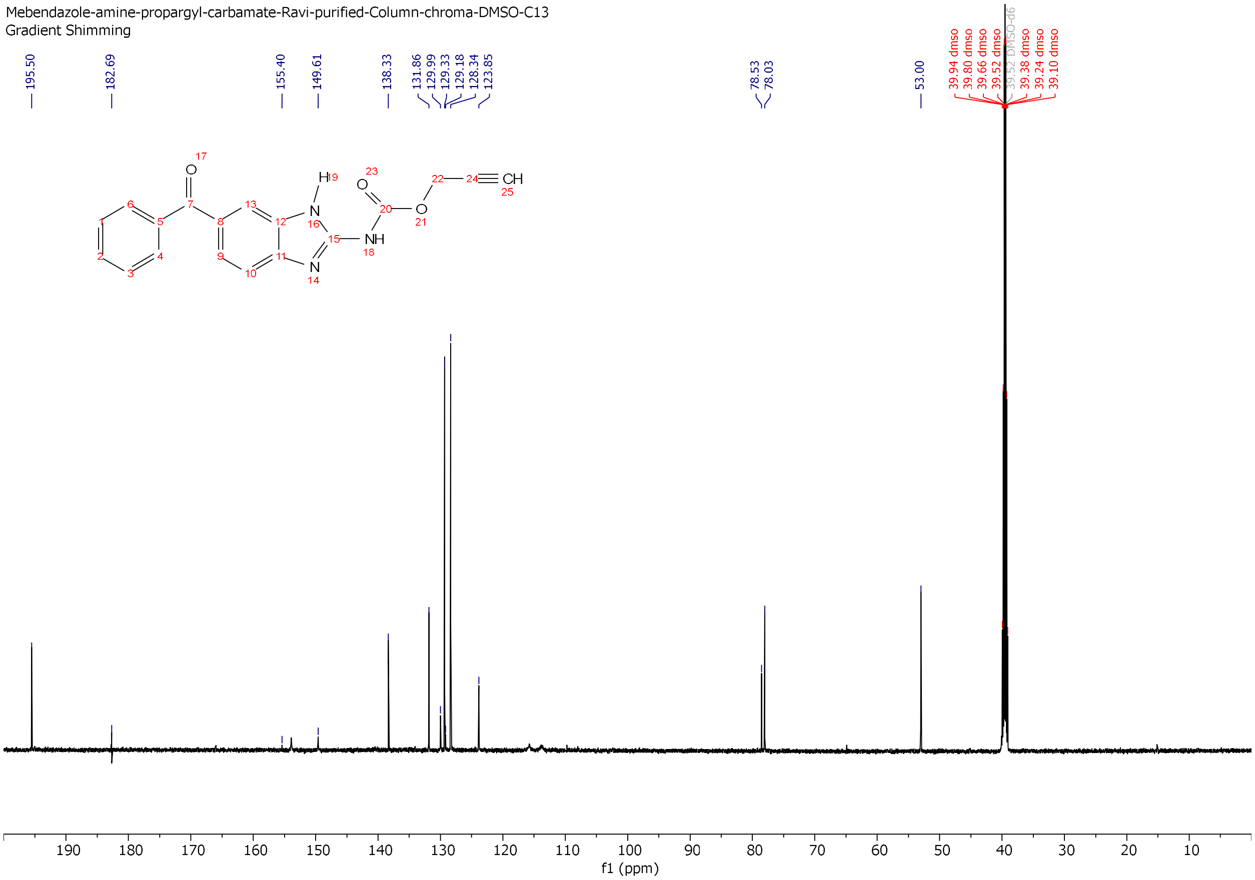

**Figure**: $^{13}$C-NMR of Prop-2-yn-1-yl (6-(benzene-2-carbonyl)-1H-benzo[d]imidazol-2-yl) carbamate

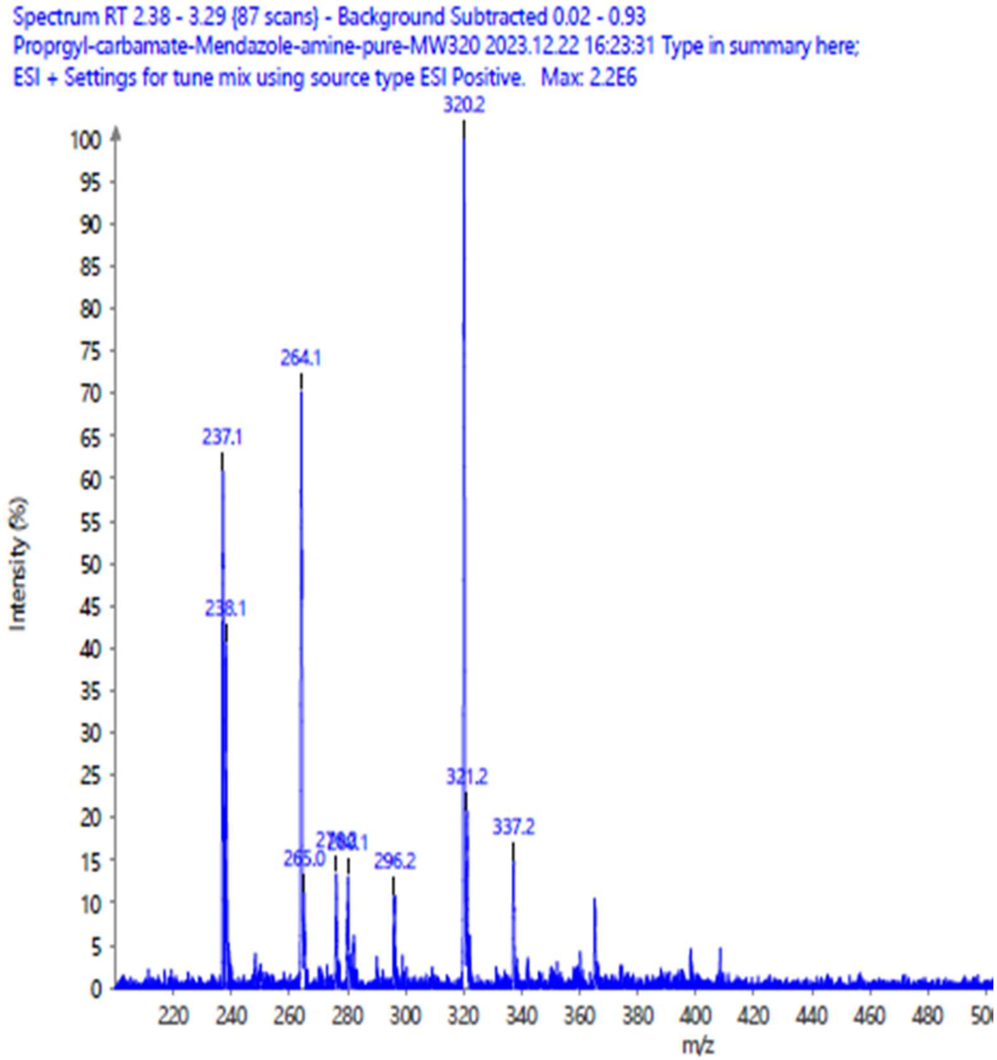

**Figure**: ESI Mass spectrum of Prop-2-yn-1-yl (6-(benzene-2-carbonyl)-1H-benzo[d]imidazol-2-yl) carbamate

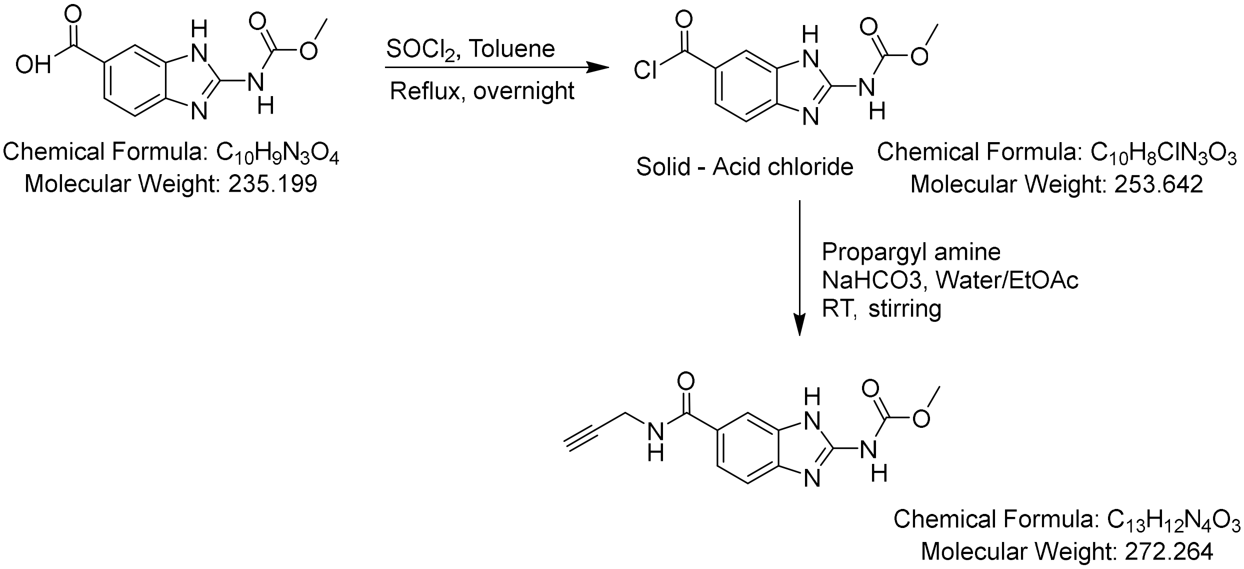

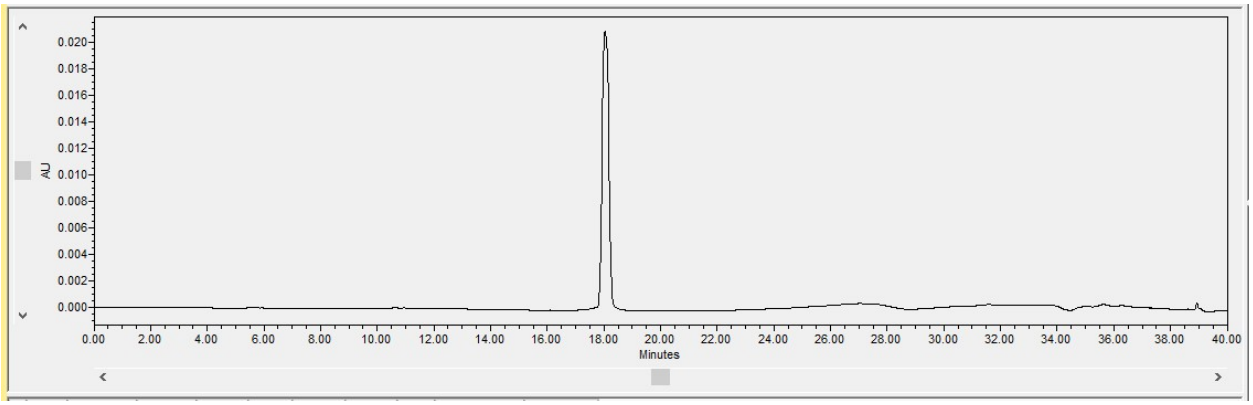

Analytical HPLC chromatogram for Prop-2-yn-1-yl (6-(benzene-2-carbonyl)-1H-benzo[d]imidazol-2-yl) carbamate

**Carboxy-propargylamide-bendazole synthesis**:

To a solution of 2-((methoxycarbonyl)amino)-1H-benzo[d]imidazole-6-carboxylic acid (235.0mg, 1.0mmol) in anhydrous toluene (20 mL) was added thionyl chloride (300 uL, excess) under nitrogen and mixture was refluxed on an oil bath overnight. The volatiles and solvents were removed under reduced pressure using a rotary evaporator. The left-over residue was used as is without further purification. The left-over residue was dissolved in ethyl acetate: water (20 mL, 3:1) and $NaHCO_3$ (100 mg, 1.2 mmol), propargyl amine (100 mg, 1.8mmol) were added to this biphasic mixture and further stirred at room temperature. Tlc analysis indicated formation of a new compound. The reaction mixture was then transferred to a separatory funnel, aqueous layer separated. The organic layer washed with water and finally with brine, dried over anhydrous $Na_2SO_4$ and concentrated under reduced pressure to yield crude material. Silica gel chromatography of crude using hexane:ethyl acetate gradient (3:1 to 1:4) afforded pure product (201 mg, 74%). $^1$H-NMR (DMSO-$d_6$) δ 3.09 (t, *J*=1.0Hz, 1H, CH), 3.77 (s, 3H, OMe), 4.05 (d, *J* =1Hz, 2H, $CH_2$), 7.42 (d, *J* =6Hz, 1H, ArH), 7.62 (dd, *J*=1 and 6 Hz, 1H, ArH), 7.94 (s, 1H, ArH), 8.79 (t, *J* =2Hz 1H, ArH), 11.74 (brs, 1H, NH). Partial $^{13}$C-NMR (DMSO-$d_6$) δ 28.5, 52.5, 72.5, 81.7, 120.5, 126.9, 148.6, 154.6, 166.5. ESI-MS: calculated *m/z* $[M+H]^+$ 273.1, observed *m/z* 273.1.

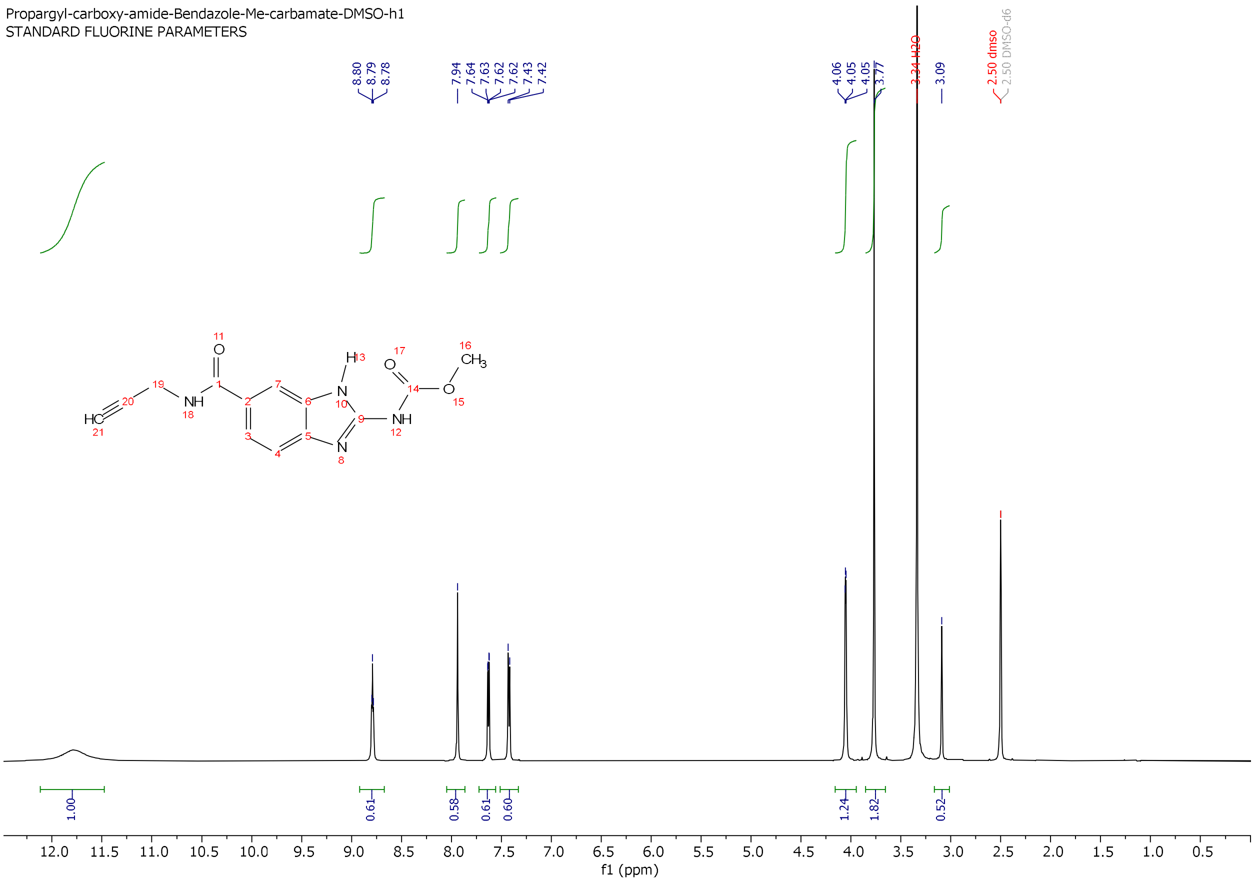

**Figure**: $^{1}$H NMR of 2-((methoxycarbonyl)amino)-1H-benzo[d]imidazole-6-carboxypropynyl amide

**Synthesis of N-(6-(phenylthio)-1H-benzo[d]imidazol-2-yl)propionamide**:

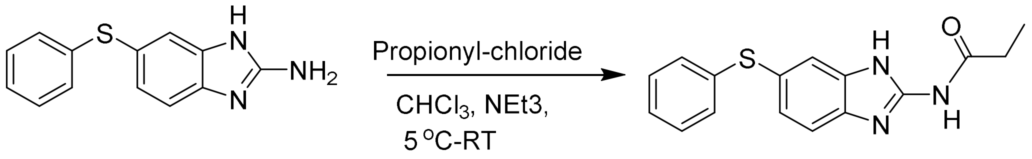

6-(phenylthio)-1H-benzo[d]imidazol-2-amine, synthesized from Fenbendazole by KOH hydrolysis[4] was used for synthesis of propionamide derivative. (241.0 mg, 1mmol) amine was dissolved in 25 mL of dry $CHCl_3$, to the clear solution was added triethylamine (305 mg, 3 mmol, 3 eq.), in a round bottom flask equipped with dry nitrogen gas filled ballon, which was placed in ice water bath 5 $^{o}$C. Propionyl chloride (95.0mg, 1.03mmol, 1.0 eq) was added slowly via syringe, the mixture was stirred 30 min. in ice bath, allowed to warm to room temperature. 10 mL dil. HCl (0.1M) solution was added to the reaction mixture, organic layer separated washed with dil. $NaHCO_3$ and finally with brine. The organic layer was dried over $Na_2SO_4$ and concentrated under reduced pressure using rotary evaporator. The dried solid residue was purified on silica column using hexanes:chloroform gradient. Fractions containing desired product as analyzed by ESI analysis were combined and concentrated to afford pale yellow solid (227 mg, 76%).

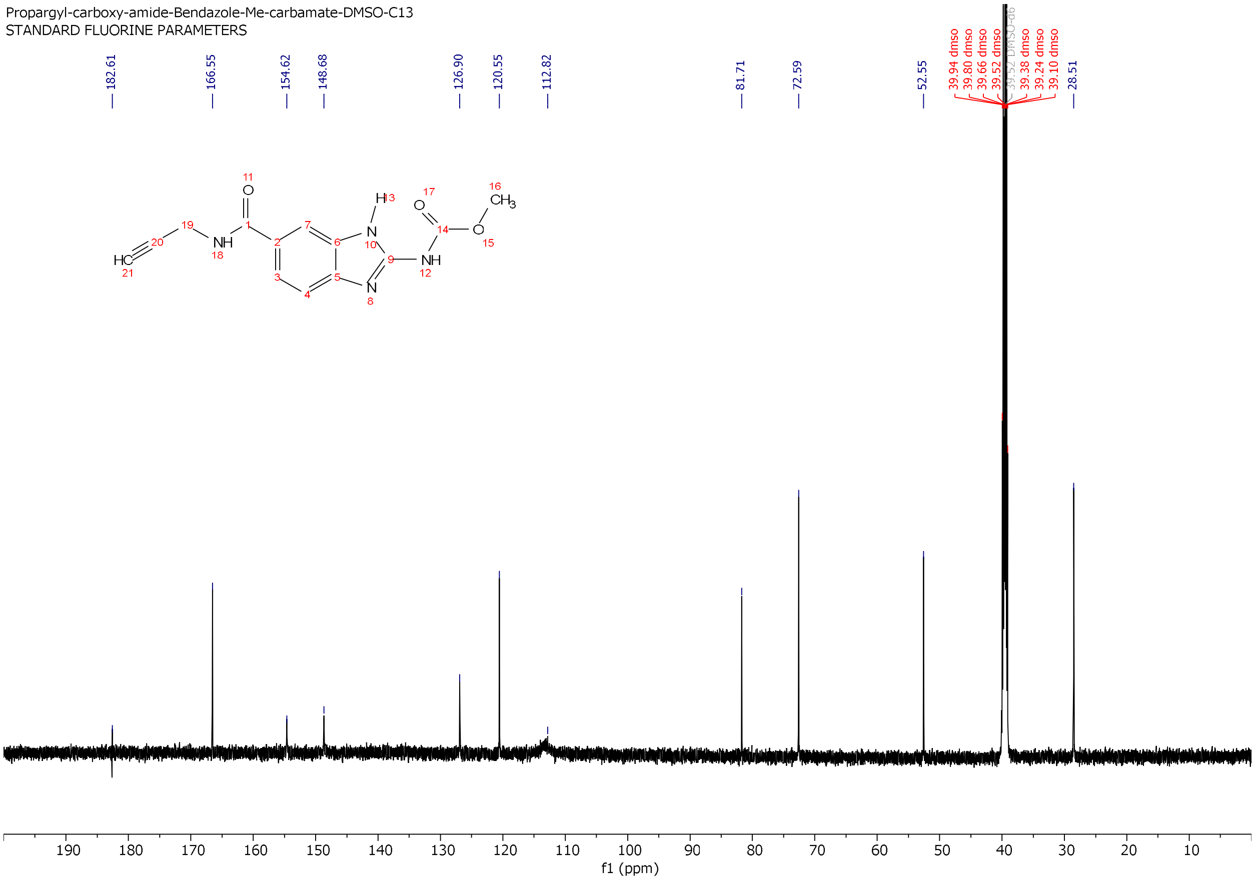

**Figure**: $^{13}C$ NMR of 2-((methoxycarbonyl)amino)-1H-benzo[d]imidazole-6-carboxypropynyl amide

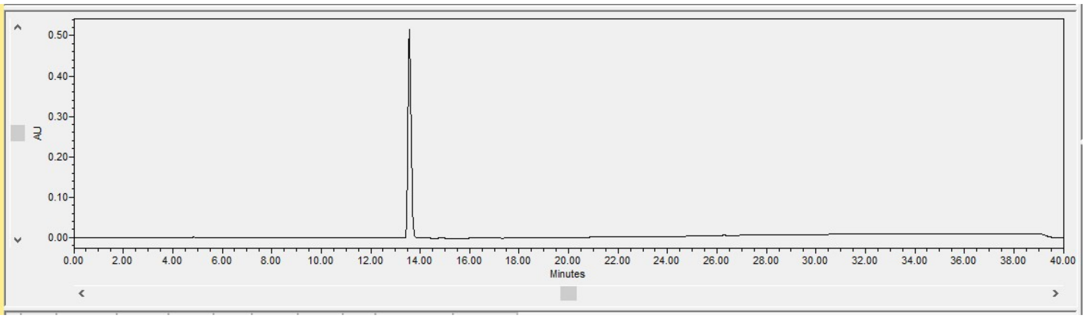

Figure: Analytical HPLC chromatogram of 2-((methoxycarbonyl)amino)-1H-benzo[d]imidazole-6-carboxypropynyl amide

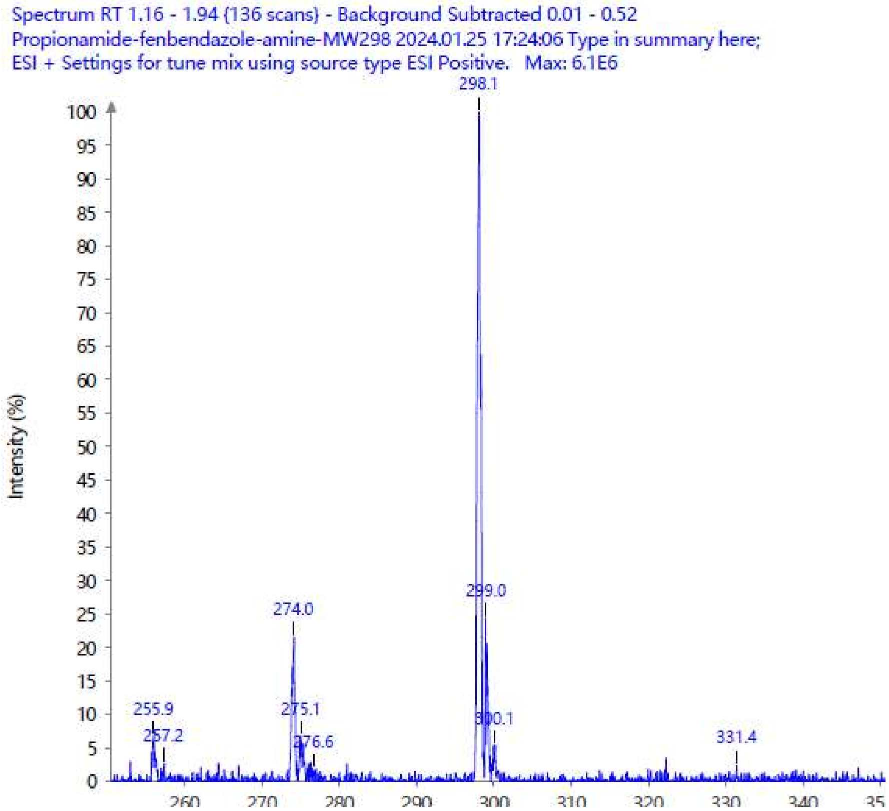

**Figure**: ESI mass spectrum of N-(6-(phenylthio)-1H-benzo[d]imidazol-2-yl)propionamide
